# Identification and characterization of nanobodies acting as molecular chaperones for glucocerebrosidase through a novel allosteric mechanism

**DOI:** 10.1101/2024.03.25.586126

**Authors:** Thomas Dal Maso, Chiara Sinisgalli, Gianluca Zilio, Isabella Tessari, Els Pardon, Jan Steyaert, Steven Ballet, Elisa Greggio, Wim Versées, Nicoletta Plotegher

## Abstract

The enzyme glucocerebrosidase (GCase) catalyses the hydrolysis of glucosylceramide to glucose and ceramide within lysosomes. Homozygous or compound heterozygous mutations in the GCase-sencoding *GBA1* gene cause the lysosomal storage disorder Gaucher disease, while heterozygous mutations are the most frequent genetic risk factor for Parkinson’s disease. These mutations commonly affect GCase stability, trafficking or activity. Here, we report the development and characterization of nanobodies (Nbs) targeting and acting as chaperones for GCase. We identified several Nb families that bind with nanomolar affinity to GCase. Based on biochemical characterization, we grouped the Nbs in two classes: Nbs that improve the activity of the enzyme and Nbs that increase GCase stability *in vitro*. A selection of the most promising Nbs was shown to improve GCase function in cell models and positively impact the activity of the N370S mutant GCase. These results lay the foundation for the development of new therapeutic routes.

## Introduction

The lysosomal enzyme glucocerebrosidase or acid-β-glucosidase (GCase, <u>EC3.2.1.45</u>) is responsible for the hydrolysis of the sphingolipid glycosylceramide (GlcCer), as well as for the transglucosylation of cholesterol by transferring the glucose moiety from GlcCer to cholesterol^1^. Biallelic mutations in the GCase-encoding *GBA1* gene cause Gaucher Disease (GD), the most common lysosomal storage disorder^2^, while heterozygous mutations in *GBA1* are the most common genetic risk factor for Parkinson’s disease (PD)^3^.

GD is characterized by a very broad spectrum of phenotypes, ranging from asymptomatic GD to patients showing hepatosplenomegaly, joint and bone pain and damage, anaemia and thrombocytopenia. Some forms of GD are neuropathic, and, depending on the neurological manifestation of the disease, GD is classified into three clinical subtypes (Type 1, Type 2 and Type 3)^4^. Type 1 GD is the most common variant and considered to be non-neurological: symptoms can be absent or systemic, and patients can develop the disease at any age. Type 2 presents the most severe neuropathic phenotype, with the onset of the disease in the first months of life and a rapid progression that leads to death during infancy. Type 3 is the chronic neuropathic form of GD, with patients surviving longer but presenting neurological symptoms over the whole course of their lives^5^. GD patients present a higher risk of developing PD, suggesting that even Type 1 GD can have neurological manifestations as patients age^6^.

PD is the second most common neurodegenerative disorder after Alzheimer’s disease, and is characterized by typical motor symptoms, such as tremor, rigidity and bradykinesia. Non-motor symptoms also occur, such as gastrointestinal defects at early stages, and psychosis and dementia at the later stages^7^. About 10% of PD cases are inherited and caused by mutations in different genes, either in an autosomal dominant or in a recessive manner. The other 90% of the cases are sporadic and can be associated with environmental exposure to toxins and pesticides and/or mutations in genes that are risk factors for the disease^7^, such as *GBA1*. GD frequency in the general population is about 1 in 50000 to 100000 (https://medlineplus.gov/), while around 10 million people worldwide are affected by PD. About 5-8% of PD patients carry a *GBA1* mutation, compared to less than 1% of healthy people. *GBA1*-associated PD presents earlier onset compared to typical sporadic PD^8,9^, and *GBA1*-PD patients are more likely to develop dementia and to die earlier compared to non-carriers^10^. More than 300 mutations in *GBA1* have been associated with GD and PD (summarized in Pokorna *et al.* 2023^11^), including nonsense mutations, deletions, insertions and missense mutations. Most of these mutations cause the GCase enzyme to be less stable and/or less active, although the exact link between the effect of the mutations and the severity of the disease is still largely unclear.

In normal conditions, GCase is synthesized by ribosomes that are bound to the endoplasmic reticulum (ER) and then transferred to the ER for the correct folding. GCase is subsequently trafficked to the lysosomes through the Golgi, while undergoing different glycosylations^12^. The two main suggested mechanisms of toxicity in GD and PD are thus linked to (i) impaired trafficking of the mutant protein (e.g. N370S or L444P) due to defective folding and stability, leading to ER stress and damage^13–15^, and (ii) reduced GCase activity, which causes the accumulation of substrates within the lysosomes and lysosomal dysfunction^2^. In neuronal cells, the impaired lysosomal function due to GCase mutation is associated with the aggregation of alpha-synuclein, with mitochondrial dysfunction and defective calcium handling^16,17^. Interestingly, GCase activity is reduced in sporadic PD brains^18^, even without accumulation of GCase substrates^19^, indicating a multilevel involvement of the enzyme in PD.

Currently, no cure for either GD or PD is available. GD patients are commonly treated by one of two strategies. A first strategy is enzyme replacement therapy (ERT), where recombinant GCase is provided intravenously. This strategy typically restores systemic symptoms but is unable to affect the neuropathic ones because of the inability of the recombinant enzyme to cross the blood-brain barrier (BBB). A second possible but more limited approach is substrate reduction therapy (SRT), through inhibition of glucosylceramide synthase, thus reducing the accumulation of GCase substrates by hampering their rate of biosynthesis. For *GBA1*-associated PD, a brain-permeant glucosylceramide synthase inhibitor, termed venglustat, was evaluated in clinical trials, but with no success in ameliorating the disease phenotype^20^.

Other possible therapeutic strategies targeting GCase both in PD and GD have been put forward, with the aim of stabilizing or activating the protein, or improving its trafficking to the lysosomes. Initially, this “molecular chaperone” approach was attempted unsuccessfully by exploiting iminosugar-based inhibitors, such as isofagomine^21^, to stabilize GCase by binding to the active site of the enzyme. High throughput screening also allowed to identify non-iminosugar inhibitors able to act as chaperones for GCase when studied *in vitro* on the recombinant enzyme and on spleen lysates obtained from GD-patients carrying the N370S *GBA1* mutation^22^. The quinazoline modulator JZ-4109 was shown to stabilize wild-type and N370S mutant GCase, and increase GCase abundance in PD- and GD-derived fibroblast cells^23^. Nevertheless, also JZ-4109 inhibits GCase activity by binding very close to the active site of the enzyme, suggesting it needs to dissociate once the enzyme reaches the lysosomes to avoid competition with the substrate. No information of further development of the molecule and (pre-)clinical applications are available to our knowledge. More recently also non-inhibitory small molecules acting as molecular chaperones for GCase were discovered, and their positive impact on iPSC-derived neurons from PD patients was demonstrated^24,25^. Other molecules that were shown to improve GCase function, and are currently in clinical trial for PD, are ambroxol and BIA-28-6156 / LTI-291^26–28^, while their mechanism of action is still largely unclear. Although their exact binding mode is unknown, docking studies suggest that also these molecules might bind close to the GCase active site ^29,30^. In the current study, we explored and validated the use of nanobodies (Nbs) as chaperones to improve (mutant) GCase stability, trafficking and activity. Nbs are small (±15 kDa) and stable single-domain fragments derived from camelid heavy chain-only antibodies^31^. Owing to their small size, stability and ease of cloning and recombinant production, Nbs have frequently been used as tools in research, in diagnostics and even as therapeutics^31,32^. They have the tendency to insert themselves into clefts or cavities on the surface of their antigens^33^, thus stabilizing certain protein conformations and/or modifying enzyme activity^34–38^. As such, Nb-mediated stabilization of GCase appears as an appealing approach to improve GCase folding in the ER and its transport to the lysosome, thus possibly reducing the amount of unfolded mutant GCase in the ER and increasing the amount of functional GCase in the lysosome, or even increase the enzymatic activity of GCase in the lysosome *per se*. With this strategy in mind, we were able to identify and characterize different sets of Nbs that stabilize or activate GCase *in vitro* and in cell models, using an allosteric mechanism that differs, to the best of our knowledge, from the currently available GCase chaperones.

## Results

### Identification of GCase-targeting nanobodies

To generate nanobodies (Nbs) specifically binding human lysosomal glucocerebrosidase (GCase), a llama was immunized with a commercial source of GCase (Velaglucerase, VPRIV®)^39–41^. To maximize the chances of obtaining a large repertoire of Nbs, which preferentially bind irrespective of the glycosylation pattern of GCase and outside its active site pocket, different phage display selection strategies were used in parallel. Hereto, GCase was first deglycosylated with Peptide:N-glycosidase F ( PNGase F) and/or allowed to react with the covalent inhibitor conduritol-β-epoxide (CBE)^42^, resulting in the following combinations that were used for two subsequent rounds of phage display panning: (i) glycosylated GCase, (ii) glycosylated GCase bound to CBE, (iii), deglycosylated GCase, (iv) deglycosylated GCase bound to CBE. Moreover, to ensure that the selected Nbs also bind at the low lysosomal pH, washing steps using a buffer at pH 5.4 were incorporated in the phage display protocol. Sequencing of the Nb open reading frames resulting from these 4 selection strategies provided 38 unique sequences. These can be grouped into 20 sequence families, where members within a sequence family display > 80% sequence identity in their complementary determining region 3 (CDR3). One representative of each family was recloned in a pHEN29 vector and subsequently expressed in the periplasm of *E. coli* as a C-terminally LPETGG-His6-EPEA-tagged protein, and the corresponding 20 Nbs were purified to homogeneity (**Supplementary Figure S1**).

### The Nbs bind GCase mostly in a glycosylation-independent way and with a variety of affinities

The binding of the purified Nbs to GCase (Velaglucerase) was first confirmed using ELISA, with binding defined as an ELISA signal of at least 3-fold above the GCase background and 3-fold above the signal of an irrelevant Nb. Using that criterion, we find binding for 11 out of the 20 tested Nbs (Nb1, Nb2, Nb3, Nb4, Nb6, Nb7, Nb8, Nb9, Nb10, Nb16, Nb17) (**Figure 1A**). Interestingly, while no binding signal is observed for Nb12 and Nb18 on glycosylated GCase, a clear binding signal is obtained when using deglycosylated GCase (**Supplementary Figure S2**). Also, for Nb10 a stronger ELISA signal is observed when using deglycosylated GCase. This is in good agreement with the fact that these Nbs result from phage display panning on deglycosylated GCase.

**Figure 1:**
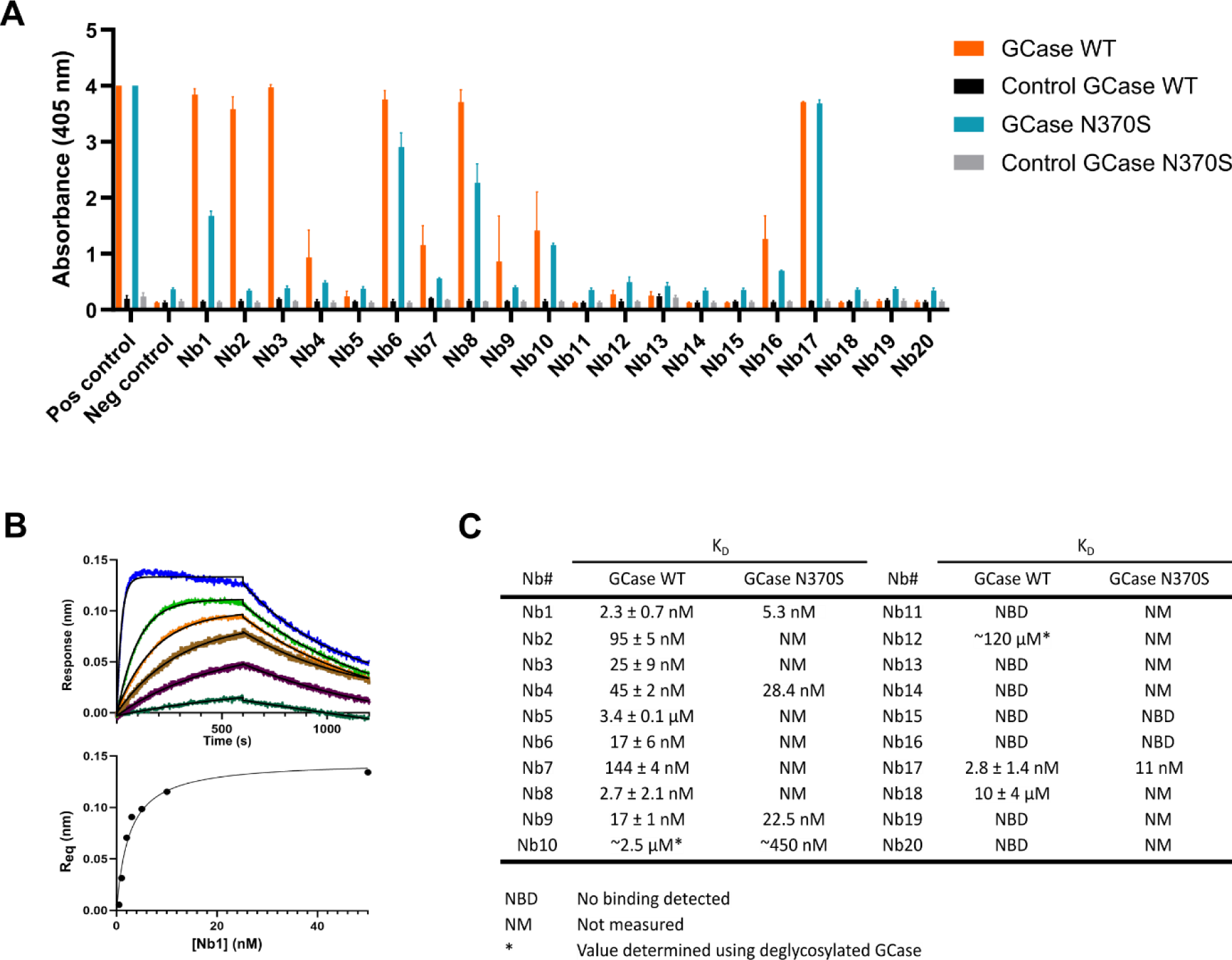
Binding of the set of 20 nanobodies to wild-type and N370S GCase. **(A)** ELISA of 20 purified Nbs using GCase or N370S GCase coated on the bottom of the ELISA wells. An irrelevant Nb is used as negative control, while the positive control displays the signal of a Nb (Nb17) directly coated in the ELISA plate. Each ELISA signal is the result of three independent experiments represented as mean values (bars) with standard deviations (error bars). **(B)** Representative Bio-Layer interferometry (BLI) traces for binding of Nb1 to wild-type GCase (upper panel) and fitting of the signal amplitudes (Req) on the Langmuir equation to determine the KD values (lower panel). **(C)** Equilibrium dissociation constant (KD) values of the full set of Nbs for GCase or N370S GCase. Values are determined by fitting of the signal amplitudes (Req) on the Langmuir equation. KD values for GCase WT are represented as mean values ± standard deviations (n = 3), while reported KD values for GCase N370S are the result of a single experiment.

To determine the binding affinities (KD) of the Nbs for (glycosylated) GCase, we next turned to biolayer interferometry (BLI). Hereto, randomly biotinylated GCase was immobilized on a streptavidin biosensor and titrated with increasing amounts of each of the Nbs, after which the equilibrium signals were plotted against the Nb concentrations and fitted on a Langmuir equation (**Figure 1B and 1C; Supplementary Figure S3**). Overall, the obtained KD values are in good agreement with the trend observed in ELISA. KD values in the sub-micromolar range are obtained for 9 of the Nbs, with Nb1, Nb8 and Nb17 showing the highest affinities (KD < 5 nM). Nb5 and Nb18 show low-affinity binding (KD in µM-range), while binding of Nb10 and Nb12 could only be observed when using deglycosylated GCase. No clear binding signals in BLI were obtained with Nb11, Nb13, Nb14, Nb15, Nb16, Nb19 and Nb20. Remarkably, while no binding was observed in BLI for Nb16, a clear binding signal was present in ELISA.

### Different subsets of Nbs affect the activity and thermal stability of GCase *in vitro*

To characterize the impact of the 20 Nbs on the activity of wild type GCase, we took advantage of a 4-MU based assay, as previously reported^43^. This approach allowed us to evaluate if any of the Nbs were able to increase the GCase activity *in vitro*, either by slowing down the time-dependent unfolding of the enzyme or by allosterically activating it. Hereto, GCase (Velaglucerase) was incubated for 30 minutes at 37°C in the presence of each Nb to allow GCase-Nb complex formation (if any), then the 4-MU substrate was added and the mix was incubated for 90 minutes before the reaction was stopped to proceed with the measurement (**Figure 2A**). This allowed to identify 7 Nbs able to significantly increase the activity of the wild type GCase: Nb3, Nb4, Nb5, Nb6, Nb10, Nb16 and Nb18. Among them, 5 Nbs were able to augment the GCase activity by more than 2-fold (Nb3, Nb5, Nb6, Nb16, Nb18), while the increment due to Nb4 and Nb10 was between 1.5 and 2-fold. Interestingly, also Nb1, Nb2, Nb8, Nb9 and Nb19 showed a trend toward a positive impact on the enzymatic activity of the GCase (between 1.5 and 2-fold), albeit not statistically significant, while the others seem to have no effects on the enzyme activity (**Figure 2B**). Isofagomine was used as a control and proved to reduce the GCase activity by about 70%. To understand if the presence of other proteins or cofactors would change the outcome of these measurements, we performed the 4-MU activity assay on cell lysates expressing wild type GCase, following the same experimental setup. Interestingly, in this case only Nb10 and Nb16 showed a significant > 2-fold increase in GCase activity, while Nb9 and Nb18 were able to significantly improve the activity by 1.5 to 2-fold. Under these conditions, some Nbs were also showing a mild decrease in GCase functionality, i.e. Nb2, Nb6, Nb17 and Nb19 (**Figure 2C**), even though not significant. This suggested that different experimental conditions may impact on the interaction between the different Nbs and the enzyme, affecting the improvement of the enzymatic activity *in vitro*. Overall, these results led us to conclude that Nb9, Nb10, Nb16 and Nb18 increase the GCase activity under both tested conditions.

**Figure 2:**
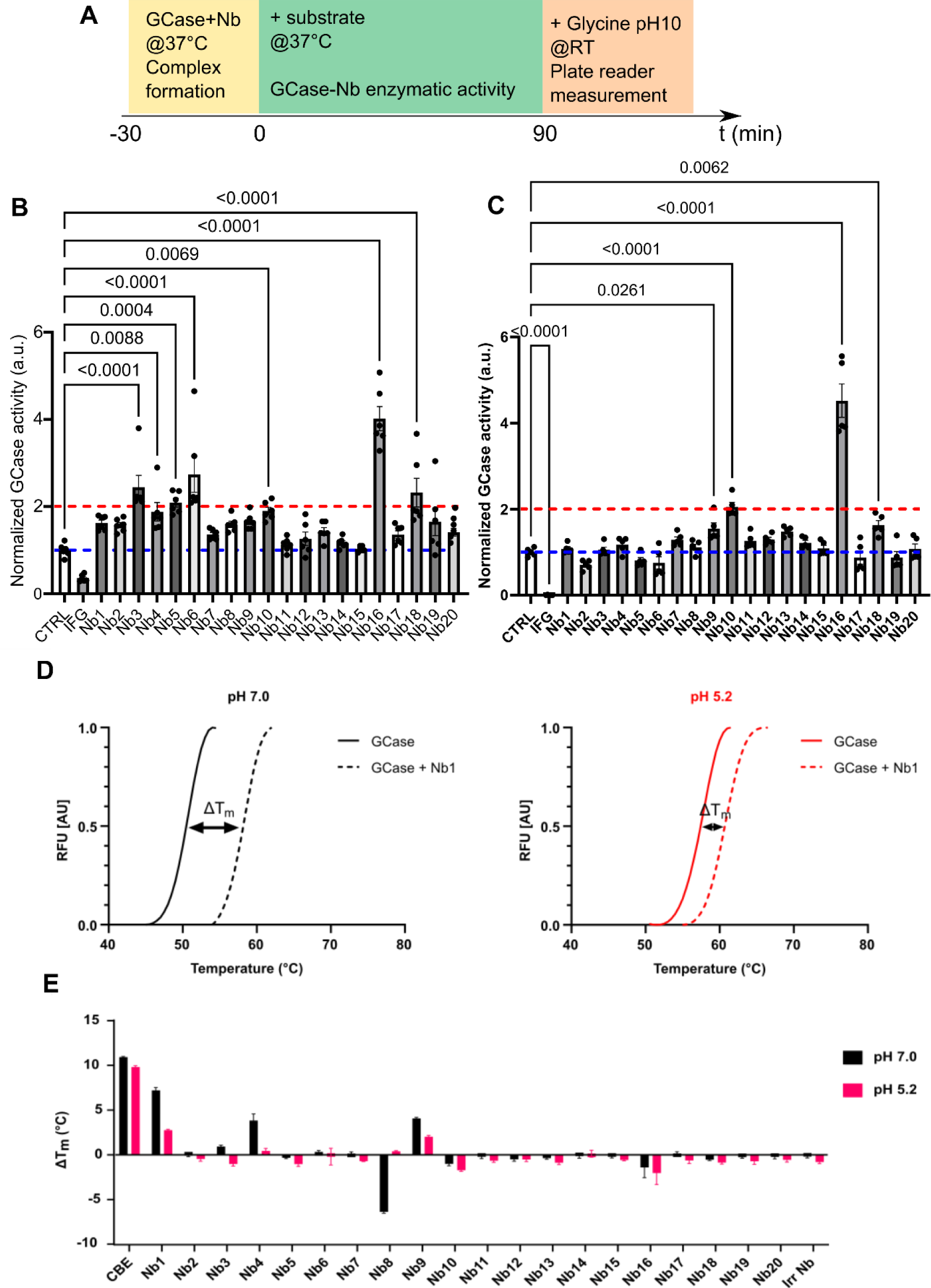
*In vitro* effect of the Nbs on GCase activity and thermal stability (Tm) **(A)** Schematic representation of the protocol for the *in vitro* 4-MU GCase activity assay; **(B)** Velaglucerase activity in the presence of each Nb of the set showed that a subset of Nbs is able to significantly improve *in vitro* GCase enzymatic activity. Isofagomine (IFG) was used as a negative control (n=6 in three independent experiments, data represented as mean±SEM, statistical analysis was performed using an Ordinary One Way Anova multiple comparison test, DF Nbs = 21, DF residual = 110, F value = 18.08); **(C)** GCase activity assay in cellular lysates in the presence of each Nb of the set showed that a subset of Nbs is able to significantly improve *in vitro* GCase enzymatic activity in the cellular lysates, quite coherently with the impact of Nbs on the Velaglucerase activity. Isofagomine (IFG) was used as a negative control (n=5 in three independent experiments, data represented as mean±SEM, statistical analysis was performed using an Ordinary One Way Anova multiple comparison test, DF Nbs = 21, DF residual = 88, F value = 47.54); **(D)** Thermal unfolding curves of GCase, obtained using a thermal shift assay at pH 7.0 (black line) and pH 5.2 (red line) in absence or presence of Nb1. **(E)** Overview of the results of the TSA assay for the full set of 20 Nbs. The plotted variation of Tm (ΔTm) is the difference between the Tm of GCase and GCase + Nb. The covalent inhibitor conduritol-β-epoxide (CBE) was used as positive control, and an irrelevant nanobody (Irr Nb) as negative control. Each TSA signal is the result of three independent experiments, shown as mean values (bars) with standard deviations (error bars).

Pathogenic mutations in the *GBA1* gene can also lead to a decreased cellular GCase activity by affecting the protein stability and subsequently causing unfolding in the ER. The development of molecular chaperones that increase protein stability, and thereby assist the correct trafficking through the ER toward the lysosome, is therefore regarded as a valid therapeutic strategy^44–47^. To assess the influence of the 20 Nbs on GCase thermal stability, we used a fluorescence-based thermal shift assay (TSA)^48^. First, the melting temperature (Tm) of GCase was determined at both pH 7.0 and 5.2 (non-lysosomal and lysosomal pH), yielding Tm values of 50.8°C and 58.0°C, respectively (**Figure 2D**). Screening of the GCase stability in the presence of the full set of Nbs, yielded 3 Nbs that considerably stabilize GCase at pH 7.0, with Nb1, Nb4 and Nb9 yielding an increase in Tm of 7°C, 4°C and 4°C, respectively (**Figure 2E**). These stabilizing effects were less pronounced at pH 5.2, with Nb1 and Nb9 providing an increase in Tm of 3°C and 2°C, respectively, while Nb4 does not cause an additional stabilization at this pH. In comparison, for the covalent active-site binding inhibitor CBE we find an increased stability of about 10°C, while this compound completely destroys the enzyme’s catalytic activity.

### The structure of GCase in complex with Nb1 reveals the mechanism of stabilization

Since Nb1, Nb4 and Nb9, which belong to different sequence families, can stabilize the GCase fold, we next wondered whether they bind different regions of GCase or target a similar “stability hotspot” region of the protein. To test this, we performed a BLI-based epitope mapping of these 3 Nbs, and also included Nb17 as a high affinity but non-stabilizing and non-activating Nb control. For the epitope mapping we set up a pairwise competition-binding experiment, where in turn one of the Nbs was biotinylated and immobilized on the BLI sensor, while the other Nbs were added in excess to GCase in solution to assess their effect on the binding of GCase. This experiment shows that Nb1, Nb4 and Nb9 compete with each other for GCase binding, while this is not the case for Nb17 (**Supplementary Figure S4**). This proves that all the stabilizing Nbs bind to the same or a closely overlapping epitope of GCase, probably implying that this corresponds to an important region for protein stability. In contrast, Nb17 binds to a different region of GCase.

To identify and characterize the stability hotspot in GCase, as well as the stabilizing mechanism of the Nbs, we co-crystallized GCase with Nb1. Hereto, we used a deglycosylated form of imiglucerase (Cerezyme®) as a source of GCase, which was mixed in a 1:1.2 ratio with Nb1 before setting up crystallizations^41^. Diffraction-quality crystals were obtained using 1.6 M magnesium sulfate heptahydrate and 0.1 M MES pH 6.5 as crystallization solution, and diffraction data were collected and the structure was refined to 1.7 Å resolution. The refined structure shows one molecule of GCase bound to one molecule of Nb1 in the asymmetric unit (**Figure 3A**). As previously described, the structure of GCase displays a globular fold formed by 3 non-contiguous domains: domain I (residues 1-29 and 383-414) is a small three-stranded antiparallel β-sheet, domain II (residues 30-77 and 431-497) forms an eight-stranded β-barrel, and domain III (residues 78-382 and 415-430) adopts a (β/α)8 triose-phosphate isomerase (TIM) barrel, which contains the active site and the two catalytic residues E235 and E340^39,49–51^. During refinement, sugar moieties were added on residues N19 and N270, which are well-known and characterized glycosylation sites^39^. Additionally, two magnesium ions have also been included in the GCase structure, one of which is located in the active site and directly interacts with the two catalytic glutamate residues.

**Figure 3:**
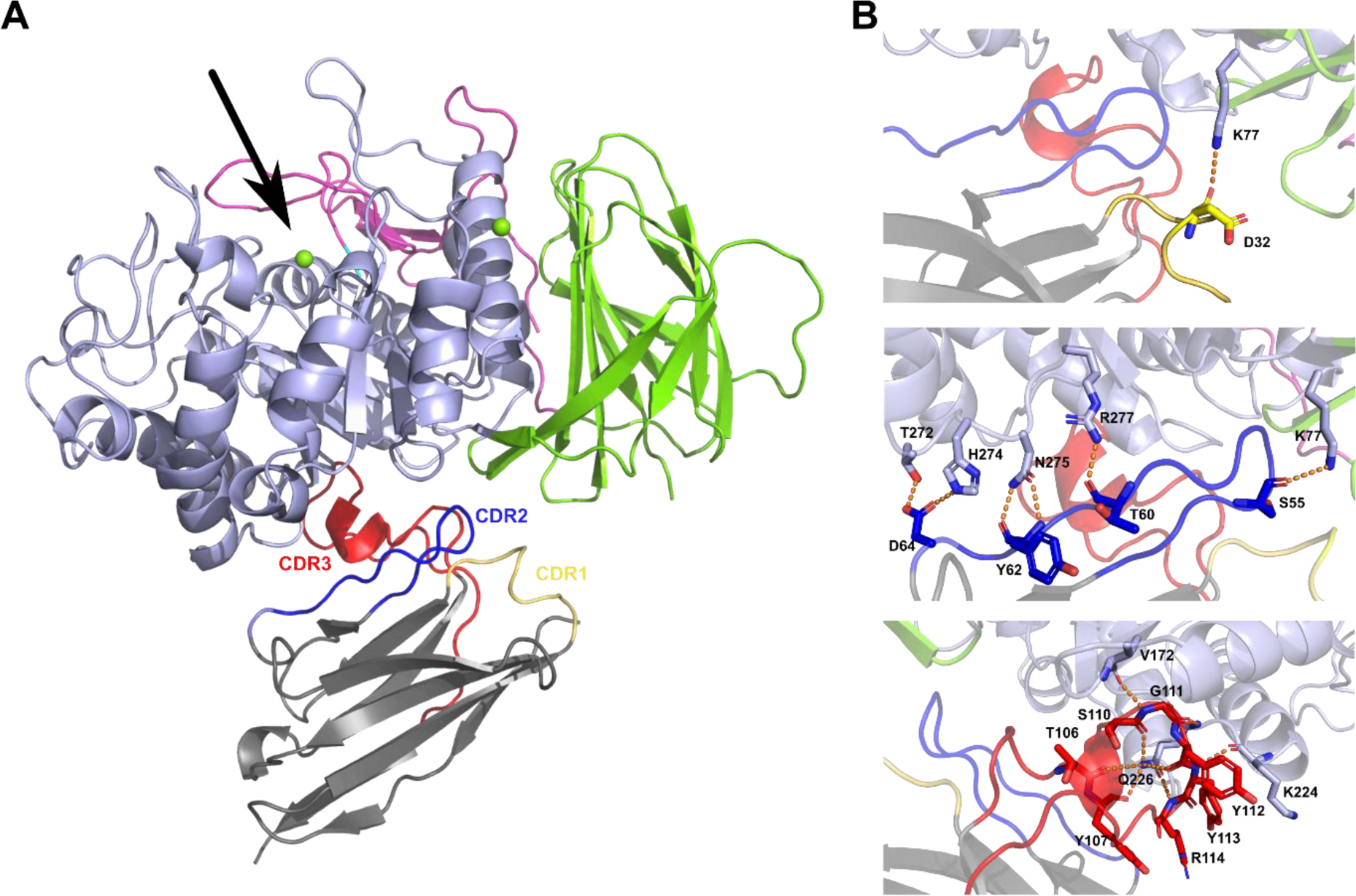
Structure of the GCase-Nb1 complex. **(A)** Overall structure of the GCase-Nb1 complex. The three domains of GCase are colored in pink (domain 1 = three-stranded anti-parallel β-sheet domain), green (domain 2 = Ig-like domain) and light blue (domain 3 = (β/α)8 triose-phosphate isomerase (TIM) barrel). Nb1 is colored grey with the CDR1 loop in yellow, CDR2 loop in dark blue and CDR3 loop in red. The position of the active site of GCase is indicated with an arrow. **(B)** Close up view of some of the interactions between the Nb1 CDR1 loop (upper panel), CDR2 loop (middle panel) and CDR3 (lower panel) with GCase. Interacting residues are shown in stick representation.

Nb1 binds to GCase on the opposite side of the active site and interacts at the interface between domain II and III (**Figure 3A**). The observation that Nb1 is binding far from the GCase active site is in good agreement with our previous finding that the Nb does not negatively interfere with the catalytic activity. On the side of Nb1, most of the interactions with GCase are provided by residues from CDR2 and CDR3 (**Figure 3B**). CDR1 provides only 2 interacting residues specifically interfacing with GCase domain II, with the main chain carbonyl of its residue Asp32 forming a H-bond with GCase residue Lys77. CDR2 interacts with residues belonging to GCase domains II and III. In particular, Ser55 forms a H-bond with Lys77 of domain II, while Thr60, Tyr61, Tyr62 and Asp64 form multiple salt bridges and/or H-bonds with residues Thr272, His274, Asn275 and Arg277 from domain III. Finally, most interactions are made by the very long CDR3 loop that partially folds into a short stretch of α-helix. Residues from CDR3 make multiple interactions with GCase domain III, with in particular the side chain of Gln226 of GCase forming multiple hydrogen bonds with main chain atoms of CDR3. This binding mode of Nb1, at the interface of two GCase domains, could explain its stabilizing effect by keeping these two domains tightly together.

Superposition of the structure of the GCase-Nb1 complex on a structure of unbound GCase (PDB 1OGS), shows no major conformational changes in GCase induced by the Nb (rmsd all atoms = 0.260 Å; rmsd Cα atoms = 0.229, using chain A of PDB 1OGS) (**Supplementary Figure S5**)^50^. Only minor conformational changes are observed in the loops surrounding the entry to the GCase active site pocket, in particular in “loop 1” (residues 311-319), “loop 2” (residues 345-349) and “loop 3” (residues 394-399). Interestingly, our structure in complex with Nb1 shows GCase in a state that has some features resembling the enzyme “active state”, with loop 1 adopting a nearly helical conformation, the side chain of residue Asp315 moving in the direction of residue Asn370, and the bulky side chains of Trp348 and Arg395 oriented away from the active site^51–53^.

### ER-targeted Nb4 and Nb9 improve lysosomal GCase activity in live wild type cells, but not the overall lysosomal proteolytic activity

Based on the *in vitro* results, studying the impact of the Nbs on either recombinant wild type GCase or on endogenous GCase from cell lysates, we curated a shortlist of priority Nbs for *in cellulo* validation. The selected Nbs (Nb1, Nb4 and Nb9) showed promise due to their capacity to either bind, activate, and/or stabilize wild type GCase, with a defined mechanism of action. We then designed a set of plasmids for mammalian expression of the Nbs in fusion with a 3xFlag tag and bicistronic co-expression with either eGFP or mCherry markers. Moreover, a targeting strategy based on established endoplasmic reticulum (ER) or lysosomal signal peptides^54,55^ was attained to either assist in the folding of GCase during ER maturation, or to enhance the stability or enzymatic activity within the lysosome, respectively. As controls, in each experiment we used an expression vector for a fragment of a non-relevant and non-functional protein (defined as Mock in the experiments) and/or untransfected control cells.

We first overexpressed the three lysosome-targeted (lyso-Nbs) and three ER-targeted Nbs (ER-Nbs) in HEK293T cells, and assessed the expression levels of each Nb by western blot (**Supplementary Figure 6A and 6B**). All ER-Nbs were expressed, though at different levels, whilst lyso-Nb1 was barely detectable compared to lyso-Nb4 and lyso-Nb9 and to the same ER-Nb (**Supplementary Figure 6C and 6D**).

Next, we set out to identify the optimal experimental system for evaluating the effect of ER- and lyso-Nbs on GCase cellular properties. We first measured GCase levels and ER/lysosome response from lysates of Nbs-transfected cells. Upon expression of ER-Nbs, the levels of GCase protein were unchanged (**Supplementary Figure S7A and S7E**). Similarly, expression of lyso-Nbs did not affect GCase protein levels (**Supplementary Figure S7B and S7F**). Ectopic expression of ER- or lyso-Nbs did not alter the overall levels of calnexin, which is induced during ER stress, or of the lysosomal protein LAMP1, respectively (**Supplementary Figure S7A-C and S7B-D**).

To examine the impact of the Nbs on GCase activity only at the lysosomes and only on Nb-expressing cells, we exploited a flow-cytometry based assay to measure lysosomal GCase activity in live cells by employing the fluorogenic substrate PFB-FDGlu (5-(pentafluorobenzoylamino) Fluorescein Di-β-D-Glucopyranoside) and co-expressing the Nbs with an mCherry reporter for flow cytometry analysis of transfected cells (**Figure 4A**). As shown in figure 4, expression of lyso-Nbs did not result in any alteration of the lysosomal activity of GCase (**Figure 4B**). In contrast, ER-Nb4 and ER-Nb9 showed a ∼15% increase in the endogenous activity of lysosomal GCase, a result that supports these Nbs as promising stabilizers of GCase, able to improve lysosomal GCase activity (**Figure 4C**).

**Figure 4:**
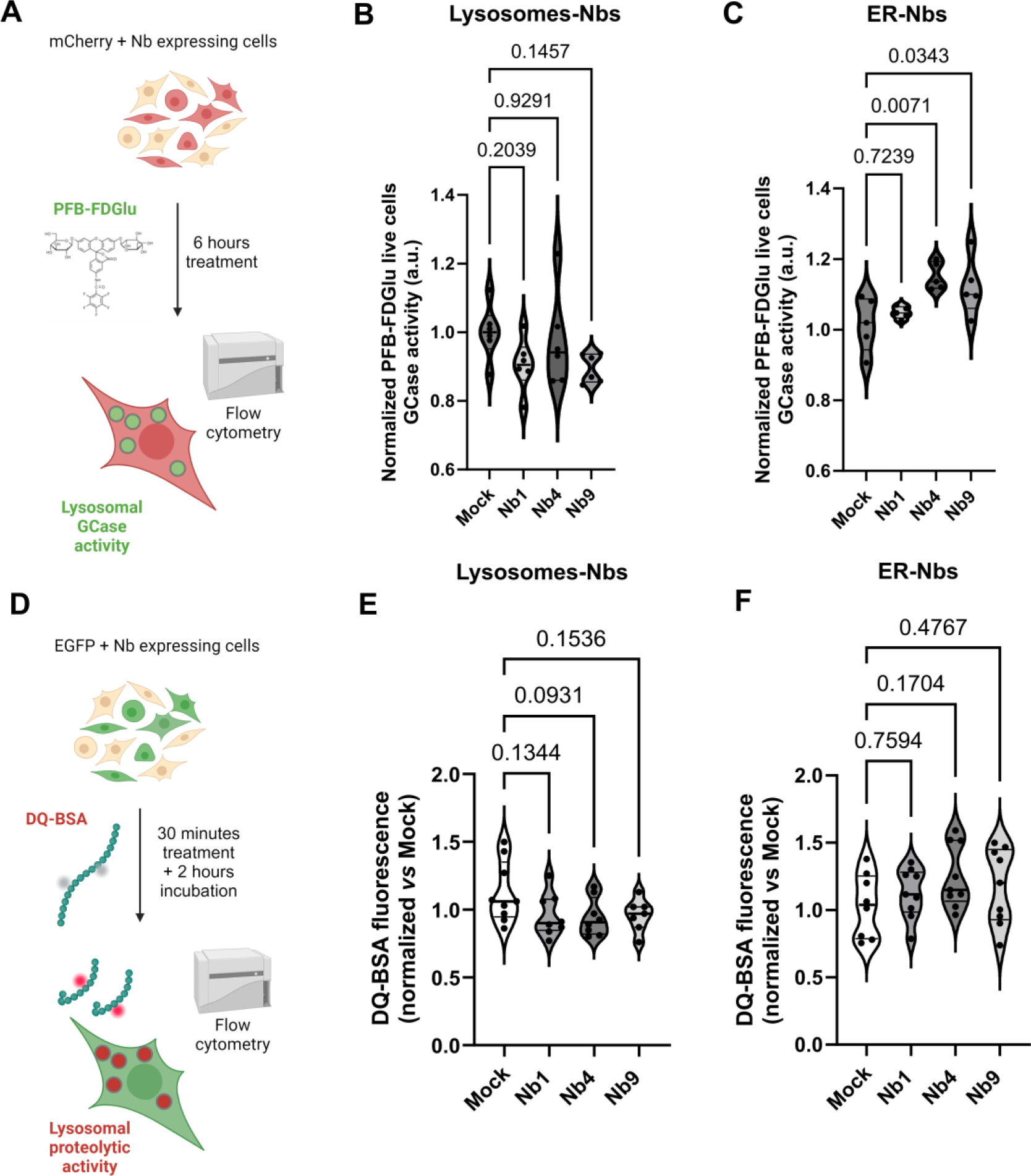
ER-targeted Nb4 and Nb9 expression increased lysosomal GCase activity in live cells, but do not improve lysosomal proteolytic activity. **(A)** Lysosomal GCase activity assay performed by using the PFB-FDGlu substrate and flow cytometry on wild type HEK293 live cells expressing Nb1, Nb4 or Nb9 targeted to the lysosomes or to the ER was performed as described in the scheme. **(B)** None of the Nbs targeted to the lysosomes show improvement in the GCase enzymatic function (n=6 in 3 independent experiments, data represented as violin plots, Shapiro-Wilk test for normality, Ordinary One Way Anova with multiple comparisons was used for the statistical analysis, DF Nbs = 3, DF residual = 20, F value = 1.909), while **(C)** expression of ER-Nb4 and ER-Nb9 provided a significant increase in the lysosomal GCase activity of about 15%, as compared to ER-Mock transfected cells (n=5 in 3 independent experiments, data represented as violin plots, Shapiro-Wilk test for normality, Ordinary One Way Anova with multiple comparisons was used for the statistical analysis, DF Nbs = 3, DF residual = 16, F value = 5.451). **(D)** DQ-BSA assay in HEK293T was performed as described in the scheme to evaluate lysosomal proteolytic activity upon Nbs overexpression. **(E)** Lysosomal proteolytic activity expressed as the ratio between DQ-BSA fluorescence mean of Nb transfected cells and Mock transfected cells (n=7-9 in three independent experiment, data represented as violin plots, Shapiro-Wilk test for normality, Ordinary One Way Anova with multiple comparisons was used for the statistical analysis, DF Nbs = 3, DF residual = 31, F value = 1.218). **(F)** Lysosomal protease activity in HEK293T cells overexpressing the selected ER-targeted Nbs (n=8-9 in three independent experiments, data represented as violin plots, Shapiro-Wilk test for normality, Ordinary One Way Anova with multiple comparisons was used for the statistical analysis, DF Nbs = 3, DF residual = 28, F value = 2.213).

Given the ability of ER-Nb4 and ER-Nb9 to improve GCase activity in live cells, we next investigated their impact on the lysosomal proteolytic activity using the DQ-Red BSA marker and a cytofluorometry-based assay^56^ (**Figure 4D**). The DQ-Red BSA dye is a fluorogenic substrate for proteases, which is hydrolyzed in the acidic compartment causing an increase in the emitted red fluorescence signal that correlates with the overall lysosomal function. As reported in figure 4E, cells transfected with the lyso-Nbs present DQ-BSA fluorescence values similar to Mock cells, suggesting that Nbs did not affect the lysosomal protease activity. Similarly, no differences were found among the different ER-targeted Nbs tested and the ER-Mock expressing cells (**Figure 4F**). This result indicates that Nbs do not impair nor enhance the overall lysosomal degradative capacity of the cell, even those able to improve GCase activity at the lysosomes. This may be explained by the fact that wild type HEK293 cells do not display any impairment in the lysosomal function that needs to be rescued.

### Selected ER-targeted Nbs improve GCase trafficking to the lysosomes

Since Nb1, Nb4 and Nb9 were shown to increase the thermal stability of GCase *in vitro* **(Figure 2E)** and Nb4 and Nb9 enhance GCase activity when targeted to the ER **(Figure 4C)**, we hypothesized that the mechanism by which the latter occurs is an ameliorated trafficking of the enzyme to the lysosomes. Thus, we assessed whether these ER-Nbs could indeed improve the trafficking of GCase to the lysosomes in cells. To this aim, the lysates of HEK293T cells overexpressing the selected ER-Nbs were incubated with ENDO H and PNGase F, two enzymes that digest glycans at different sites. Specifically, ENDO H cleaves immature glycans present in the ER but not after additional modifications that occur in the Golgi, which can be instead cleaved by PNGase F. Thus, the sensitivity of GCase to ENDO H or PNGase F is an indicator of the amount of protein present in the ER or post-ER, respectively. The higher the ratio between ENDO H-resistant and ENDO H-sensitive GCase, the more GCase is predicted to have reached the lysosomal compartment. We measured by western blot the level of total GCase and of GCase sensitive to ENDO H or to PNGase F in cell lysates (**Figure 5A**). Quantification of the ER fraction or post-ER fraction was performed following previous methods^57^. The PNGase F-sensitive GCase band was considered as a reference for the deglycosylated GCase fraction. Thus, when quantifying the different bands in the ENDO H treated GCase lane, the band that runs at the same height as the band in the PNGase F lane corresponded to the protein that is still localized at the ER. The rest of the ENDO H treated GCase bands represented the post ER GCase fraction, which was not deglycosylated by the ENDO H treatment (ENDO H resistant fraction). Quantification of the ratio between the band intensities corresponding to the post ER and to the ER GCase fraction, showed that Nb1 and Nb9 significantly increase the post ER/ER GCase fraction ratio, suggesting that they were able to promote the correct folding and trafficking of the protein through the ER and the Golgi (**Figure 5B**). Nb4 also presented a trend toward amelioration of the trafficking, even though non-significant, suggesting that the epitope the three Nbs bind is not only crucial for GCase stability but also for the trafficking of the protein. Coherently, when performing colocalization analysis between the ER marker calnexin and the ER-targeted Nbs on confocal images (**Figure 5C**), we revealed Pearson’s coefficients between 0.2 and 0.4 on average (**Figure 5D**), suggesting only partial colocalization of the Nbs at the ER. The incomplete colocalization (i.e. below 1), significantly different from the control Mock expressing cells, suggests that the ER-targeted Nb4 and Nb9 are more prone to leave the ER, likely upon binding to the GCase which is further trafficked to the lysosomes. When performing colocalization analyses between the lysosomal marker LAMP2A and the ER-targeted Nbs on confocal images (**Figure 5E**), we found Pearson’s coefficients values that are significantly increased for Nb9, up to 0.76 for certain cells, further supporting the idea that Nb9 can promote the trafficking of GCase to the lysosomes.

**Figure 5:**
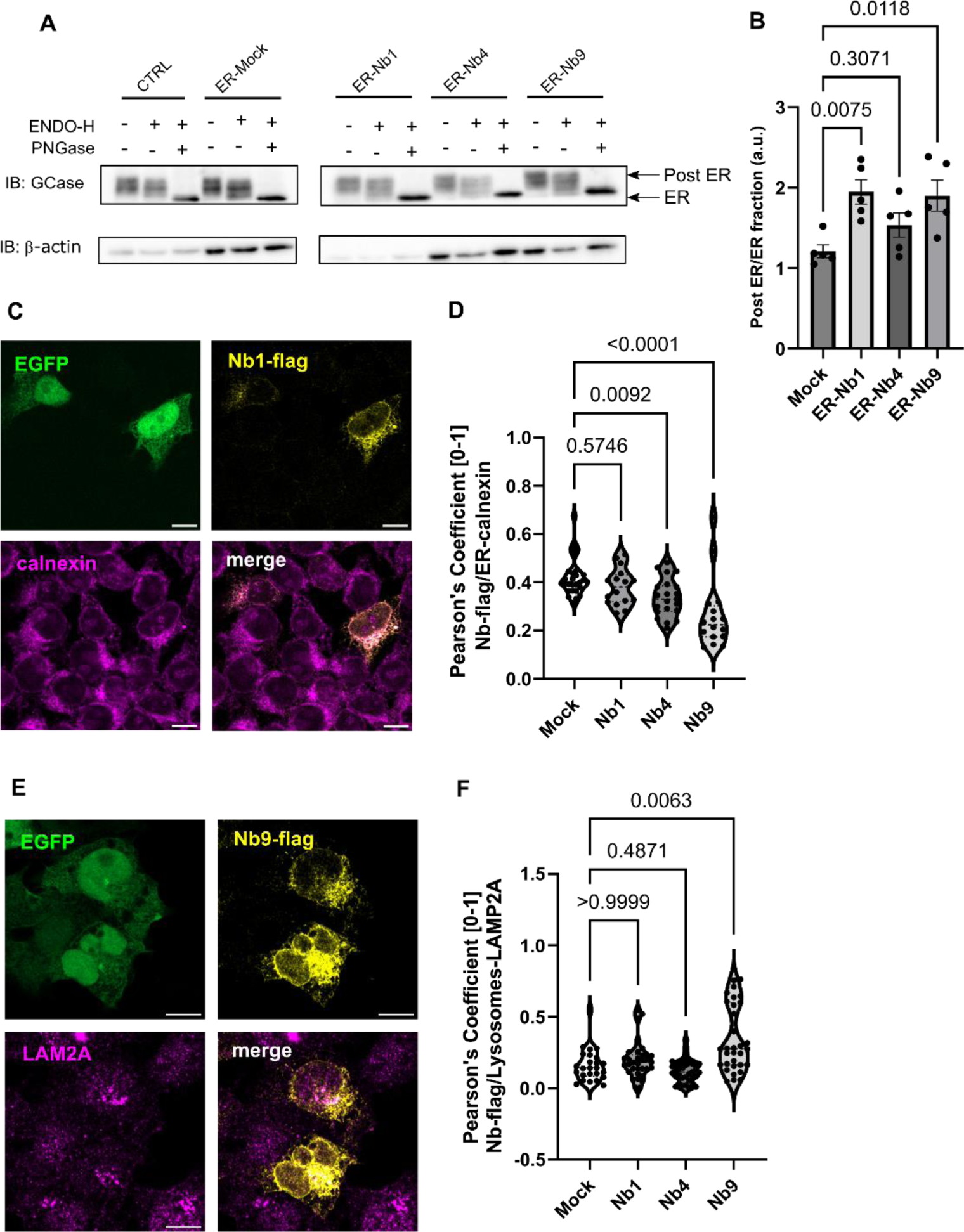
ER-Nb1, ER-Nb4 and ER-Nb9 increase GCase trafficking to the lysosomes. **(A)** Representative western blot of lysates of Nb-transfected HEK293T cells incubated with ENDO H and PNGase F to identify the ENDO H resistant (post ER, higher MW band) and the ENDO H sensitive bands (ER, lower MW band). **(B)** Ratio between the post ER and ER fraction of GCase (ENDO H resistant/ENDO H sensitive) measured in cell lysates incubated with ENDO-H and PNGase F (as a reference to detect ER fraction) show an increase for the ER-Nb1 and ER-Nb9, and a trend toward an increase for Nb4, as compared to the control (n=5, data represented as mean±SEM, Shapiro-Wilk test for Normality, Ordinary One Way Anova with multiple comparisons was used for the statistical analysis, DF Nbs = 3 DF residual = 10 F value 5.996). **(C)** Representative confocal images of HEK293T cells transfected with Nb1-flag-IRES-EGFP targeted to the ER stained for flag (yellow) and calnexin (magenta), while EGFP was used as a marker for Nb-flag expressing cells (scale bar 10 µm). **(D)** Colocalization of ER-targeted Nbs in the ER in HEK293T cells overexpressing the selected Nbs was evaluated by calculating the Pearson’s coefficient using the JACoP plugin of ImageJ between flag and calnexin staining. The results show that all the Nbs present a certain degree of colocalization with the ER-marker calnexin, which is significantly reduced in Nb4 and Nb9, as compared to the Mock transfection (n=16-28 cells in 2 independent experiments, data represented as violin plot and median values, Shapiro-Wilk test for normality, followed by Kruskal-Wallis test with multiple comparisons,). **(E)** Representative confocal images of HEK293T cells transfected with Nb9-flag-IRES-EGFP targeted to the ER stained for flag (yellow) and LAMP2A (magenta), while EGFP was used as a marker for Nb-flag expressing cells (scale bar 10 µm). **(F)** Localization of ER-targeted Nbs to the lysosomes in HEK293T cells overexpressing the selected Nbs was evaluated by calculating the Pearson’s coefficient using the JACoP plugin of ImageJ between flag and LAMP2A staining. The results show that all the Nbs present a certain degree of colocalization with the lysosomal-marker, which is significantly increased for Nb9, as compared to the Mock transfection (n=26-35 cells in 2 independent experiments, data represented as violin plot and median values, Shapiro-Wilk test for normality, followed by Kruskal-Wallis test with multiple comparisons).

### A subset of Nbs binds and increases the activity of the GCase N370S PD mutant *in vitro* and in cell models

GCase N370S is the most common mutation associated with PD and GD. This mutant shows significantly reduced catalytic activity *in vitro* and in cells^13,58^. Of note, previous studies showed that GCase is misprocessed in the ER leading to ER stress in iPSC-derived dopaminergic neurons^13^. N370S retention in the ER was also shown in PD-patient fibroblasts^59^. Thus, N370S mutant GCase is an interesting paradigm to test the potential beneficial effects of the leading Nbs. To this aim, we purified the recombinant GCase-N370S protein and tested the full set of 20 purified Nbs for binding on GCase N370S in ELISA (**Figure 1A, Supplementary Figure S8**). Nb1, Nb6, Nb8, Nb10, Nb16 and Nb17 showed a > 3-fold binding signal over the background. Nevertheless, many other Nbs that bind to wild-type GCase also gave a clear binding signal above the background for N370S, including Nb4 and Nb9.

Next, we used BLI to determine the binding affinities (KD) for a selected subset of Nbs: Nb1, Nb4, Nb9, Nb10, Nb16 and Nb17, based on the results obtained on the wild type enzyme *in vitro*, (**Supplementary Figure S9**). In this case, the Nbs were site-specifically biotinylated at their C-terminus, captured on a Streptavidin sensor and titrated with increasing concentrations of the N370S mutant. Similar to wild-type GCase, no binding was observed for Nb16. The other Nbs bind to the N370S mutant with comparable affinities to the wild-type enzyme, with the notable exception of Nb10, which showed a 5-fold higher affinity (**Figure 1C**). We next investigated the impact of the Nbs on the activity of GCase N370S *in vitro*. This experiment was performed only for a selection of Nbs that show interesting properties *in vitro* and in cells for the wild type GCase protein, using the classical 4-MU assay, as previously done for the wild type enzyme. The outcome yielded a > 2-fold significant increase in the enzyme activity for Nb10 and Nb16 (**Figure 6A**).

**Figure 6:**
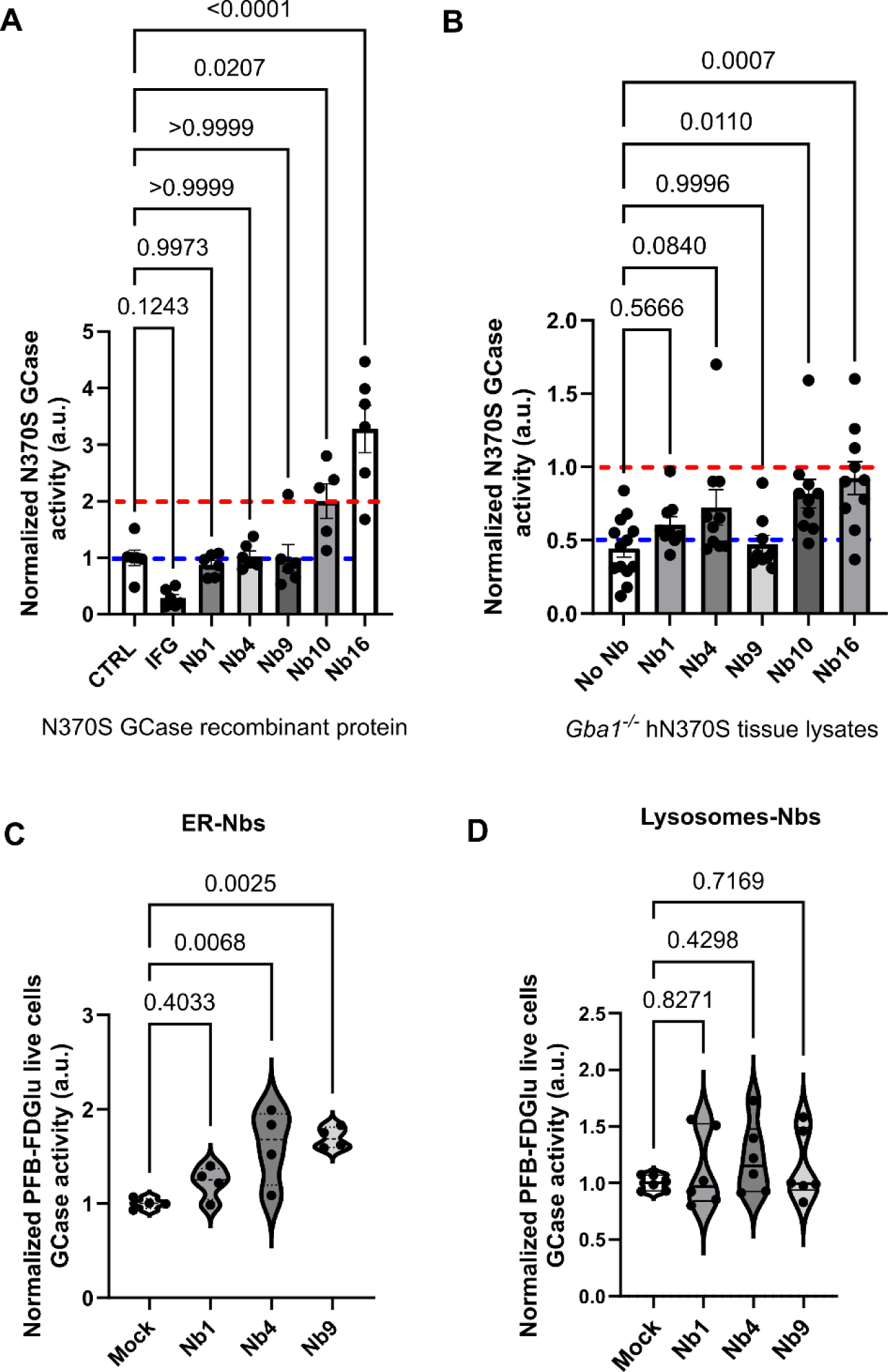
Impact of selected Nbs on the pathogenic GCase mutant N370S. **(A)** N370S GCase activity assay in the presence of Nb1, Nb4, Nb9, Nb10 and Nb16 showed that Nb10 and Nb16 are able to significantly improve *in vitro* N370S GCase enzymatic activity. Isofagomine (IFG) was used as a negative control (n=6, data represented as mean±SEM, statistical analysis was performed using an Ordinary One Way Anova multiple comparison test; **(B)** GCase activity assay in gut lysates from hN370S GCase mice in the presence of Nb1, Nb4, Nb9, Nb10 and Nb16 showed that Nb10 and Nb16 are able to significantly improve *in vitro* N370S GCase enzymatic activity. (n=4 tissue per genotype, each tested in 3 independent experiments in 3 technical replicates, data represented as violin plot showing the average value for the technical replicates for each experiment in each biological sample, statistical analysis was performed using an Ordinary One Way Anova multiple comparison test, DF Nbs = 5, DF residual = 55, F value = 4.926). **(C)** Co-expression of N370S GCase mutant and the ER-Nb4 and ER-Nb9 in *GBA1* KO cells induced an increase in the lysosomal N370S GCase activity of about 60-70%, as compared to ER-Mock transfected cells, while no effects were observed for ER-Nb1 (n=4, in two independent experiments, data represented as violin plots, an Ordinary One Way Anova test with multiple comparisons, after Shapiro-Wilk Normality test was used for the statistical analysis Df Nbs = 3, Df Residual 12, F = 8.352). **(D)** Co-expression of N370S GCase mutant and the lysosomal-targeted Nbs in *GBA1* KO cells did not show any effect on the lysosomal N370S GCase activity, as compared to lysosomes-Mock transfected cells (n=6 in 3 independent experiments, data represented as violin plots, an Ordinary One Way Anova test with multiple comparisons, after Shapiro-Wilk Normality test was used for the statistical analysis, DF Nbs = 3, DF residual = 20, F value = 0.5994).

To corroborate these results in a mouse model, we measured the effect of these Nbs on GCase activity using gut lysates from the *Gba1^-/-^ h*N370S mice, which express the human *GBA1* N370S gene in a null murine *Gba1* background^60^, and present a reduced GCase activity in vitro (**Supplementary Figure 10**). As **Figure 6B** illustrates, addition of Nb10 and Nb16 to *Gba1^-/-^ h*N370S gut lysates significantly increased GCase activity by about 2-fold, with a trend of increase also observed for Nb4. Overall, these findings indicate that Nb10 and Nb16 are efficient activator of GCase in vitro, emerge as promising candidates for correcting mutant GCase activity.

To investigate the possible impact of the stabilizing Nbs, i.e. Nb1, Nb4 and Nb9, in improving the trafficking and lysosomal activity of N370S GCase in live cells, we exploited a CRISPR/Cas9 *Gba1* knock down (KD) model (**Supplementary Figure 11**) in which we co-expressed the N370S mutant and the different Nbs. Both the clones 6E and 11F *GBA1* KD cells presented reduced GCase level by western blot (**Supplementary Figure 11A and 11B**) and reduced lysosomal GCase activity in live cells (**Supplementary Figure 11C**), as evaluated using the PFB-FDGlu substrate assay. We chose to exploit the 11F *GBA1* KD cells, which presented the larger reduction in GCase levels and activity, to overexpress the N370S GCase mutant. Upon overexpression, we verified that > 80% of the N370S expressing cells were also expressing the Nbs by immunocytochemistry (**Supplementary Figure 11D**), and that N370S overexpression induced an overall increase in lysosomal GCase activity in 11F *GBA1* KD live cells, despite the incomplete transfection efficiency (**Supplementary Figure 11E**). When overexpressing Nb4 and Nb9 targeted to the ER, the increase in the N370S GCase lysosomal activity was between 60% and 70% as compared with Mock overexpressing cells (**Figure 6C**). No effects were observed for ER-Nb1 (**Figure 6C**) and for all the lysomes-targeted Nbs (**Figure 6D**). This is in good agreement with the fact that N370S is known to present an impaired trafficking to the lysosomes, which may be rescued by the expression of the Nb4 and Nb9 at the ER.

## Discussion

The lysosomal enzyme glucocerebrosidase (GCase) plays a fundamental role in the complex cellular pathways linked to lysosome-dependent autophagy and proteostasis^13,16,61,62^. Biallelic (homozygous or compound heterozygous) mutations in the gene encoding GCase (*GBA1*) are causative for the lysosomal storage disorder GD, whereas heterozygous mutations are the most important genetic risk factor for the development of PD^3^. Importantly, GCase dysfunction was also observed in models or patients of other inherited forms of PD and in sporadic PD^18,63^. Hence, GCase is considered as an appealing therapeutic target for both disorders. Most of the disease-linked *GBA1* mutations cause defects in GCase folding, stability, trafficking and/or activity, suggesting that possible strategies to target this protein and ameliorate its function would involve the development of pharmacological chaperones able to act as stabilizers and/or activators. However, while some of such currently available chaperone molecules were tested in clinical trials, none of them actually showed a significant impact in PD/GD models or patients and reached the market so far^64^.

Here, we present the identification of different families of nanobodies (Nbs) directed against GCase, that were thoroughly characterized *in vitro* to reveal their ability to improve GCase stability and activity. One factor that might contribute to the failure in the clinic of many of the current molecular chaperones, including iminosugars, non-iminosugars and other types of molecules, is that they commonly bind into or close to the GCase active site pocket and hence act as inhibitors of its catalytic activity^64^. To overcome such potential problems, and to screen from the beginning for Nbs that bind outside the active site pocket and act in a true allosteric fashion, we devised a phage display selection procedure using both apo-GCase and GCase where the active site is blocked with the covalent inhibitor CBE. Additionally, to obtain Nbs with maximal versatility and the capacity to bind GCase in different cellular compartments and different mutant forms, we reasoned that it would be advantageous that the Nbs can bind to GCase irrespective of its glycosylation state. Hereto we performed phage display selections with either glycosylated or deglycosylated GCase (Velaglucerase).

The selection procedure finally yielded a panel of 20 different Nbs families, where one representative of each family was purified and used for further biochemical/biophysical characterization. The majority of the obtained Nbs show clear binding to GCase in ELISA and BLI, and display affinities in the low to high nanomolar range (**Figure 1**). Nevertheless, some other Nbs either show no binding or very low affinity binding, while they were picked up during phage display panning. This could be due to the fact that more GCase is directly coated to the immunosorbent plate during selections and that the selection conditions were different than in the ELISA. As we envisioned in our selection strategy, most of the Nbs bind in a glycosylation-independent way, except for Nb10, Nb12 and Nb18, which seem to bind better to the deglycosylated GCase (**Figure 1 and Supplementary Figure S2**).

*A priori* we envisioned two potential and possible overlapping functions for the GCase-targeting Nbs: (i) Nbs that increase the conformational stability of GCase and would thereby prevent unfolding and aid in the trafficking of GCase through the ER toward the lysosome, which we call **Class I Nbs**, and (ii) Nbs that increase the catalytic activity of GCase, which we call **Class II Nbs**. Detailed *in vitro* characterization of the purified Nbs allowed us to pinpoint these properties. First, a TSA experiment showed that three Nbs - Nb1, Nb4 and Nb9 - increase the melting temperature of GCase *in vitro* (**Figure 2**). GCase on its own is significantly more stable at the lysosomal pH than at neutral pH (Tm of 58°C and 50.8 °C at pH 5.2 and 7.0, respectively). All three Nbs stabilize GCase at neutral pH, with Tm shifts of 7 to 4°C, while only Nb1 and Nb9 also stabilize GCase at pH 5.2, albeit to a lesser extent. The stabilizing effect at neutral pH, corresponding to the pH in the ER, would potentially make these Nbs exquisite tools to facilitate the trafficking of (mutant) GCase through the ER and act as molecular chaperones. This categorizes **Nb1, Nb4 and Nb9 as Class I Nbs**. Interestingly, epitope mapping shows that all three class I Nbs bind to overlapping epitopes of GCase (**Supplementary Figure S4**), thereby unveiling a “hotspot” of protein stability on the GCase surface. To characterize this binding pocket, we solved the crystal structure of GCase in complex with the highest affinity Class I Nb, Nb1 (**Figure 3**). This structure shows that Nb1 mainly uses its CDR2 and CDR3 loops to interact with GCase, with CDR3 being particularly long and partially adopting an α-helical conformation. Nb1 binds to GCase at the opposite side of its active site and at the interface of GCase domain II and III (**Supplementary Figure S12**). Specifically, residues from the CDR2 and CDR3 loops of Nb1 interact with GCase domain III and the linker region between domain II and III, while the CDR1 loop makes two interactions with GCase domain II. We hypothesize that the stabilizing effect of the class I Nbs could stem exactly from this interaction on the interface of domain II and III, thus keeping both domains tightly together. Our structural analysis thus shows that the Class I Nbs bind to a true allosteric pocket, located at large distance from the GCase active site, which turns out to be crucial for GCase stability. To the best of our knowledge, this binding pocket is completely different from the binding sites of any previously described GCase chaperone^64^. The discovery of this novel stability hotspot on the GCase surface may also facilitate the future development of (small) molecules other than the Nbs targeting this region to improve GCase wild type or mutant properties. Important for the development of such molecular chaperones that guide GCase through the ER toward the lysosome, is the observation that the class I Nb epitope differs from the binding sites of LIMP-2 and Saposin-C (**Supplementary Figure S13**). These proteins function as a natural molecular chaperone for the import of GCase in the lysosome and as activator of GCase in the lysosome, respectively^12,52^. Computational as well as experimental methods have shown that both proteins bind in the vicinity of the GCase active site pocket, at binding surfaces that do not overlap with the Nb1 epitope (**Supplementary Figure S13**)^53,65^. This implies that any molecular chaperone binding to the newly discovered allosteric pocket, including the class I Nbs, would very likely not interfere with the binding of either Saposin-C or LIMP-2.

To identify Class II Nbs, next, 4-MU based GCase activity assays were performed with all 20 Nbs and using either purified GCase or cell lysates expressing recombinant GCase (**Figure 2**). These experiments revealed that **Nb3, Nb4, Nb5, Nb6, Nb9, Nb10, Nb16 and Nb18 act as Class II Nbs**, able to increase GCase activity in either or both assays. However, it should be noted that the experimental setup of the assays, where the GCase-Nb complex is incubated for 2 hours before the amount of product is measured, can result in an increased observed activity either by increasing the enzymatic activity *per se* or by preventing or slowing down the time-dependent unfolding of the enzyme. Surprisingly, Nb16 has the strongest activity-increasing effect (about a 4-fold increase for both pure GCase and GCase present in cellular lysates), even though its affinity for GCase is very low (binding is only observed in ELISA, see **Figure 1**). This result could potentially be explained by the different experimental conditions of the binding and activity assays, in particular the presence of the 4-MU substrate and detergents in the latter. This outcome thus poses interesting questions about the exact mode of action of Nb16 on GCase activity, and suggests that Nb16 may have valuable properties in a more physiological environment presenting membranous structures, co-factors and the actual GCase substrate.

Based on the *in vitro* results and considering the two identified classes of GCase-Nbs, we shortlisted three Nbs for further *in cellulo* studies. We selected Nb1 for its nM affinity and GCase-stabilizing effect, and Nb4 and Nb9 which belong to both the Class I and Class II Nbs. In order to overcome the limitation of the common inability of Nbs to cross intracellular membranes, we directly targeted the Nbs either to the ER or to the lysosomes. All Nbs were expressed when targeted to the ER, while only Nb4 and Nb9) could be detected when expressed in the lysosome (**Supplementary Figure S6**), potentially reflecting a difference in the vulnerability of the Nbs to the aggressive environment in the lysosome. To specifically assess the impact of these modifiers only in the sub-population of cells expressing the Nbs, we established robust flow cytometry-based assays where Nbs were co-expressed with mCherry, and GCase activity was tested with the fluorescent substrate PFB-FDGlu for in-cell recording. Although the used HEK293T cells express fully functional wild type GCase, and therefore lack any enzymatic or lysosomal defects, we still observed a 15% increase in lysosomal GCase activity in the presence of ER-Nb4 and ER-Nb9 (**Figure 4**). This aligns with the ability of these Nbs to enhance the fraction of GCase in post-ER compartments (**Figure 5**). Another interesting observation was that ER-targeted Nbs showed a reduced co-localization with the ER compared to a control ER-targeted mock sequence (**Figure 5**), suggesting that ER-Nbs bind GCase in the ER and then “hitchhike” together with the enzyme through the Golgi toward the lysosomes. This is also confirmed for Nb9 by an increased colocalization to the lysosomes as compared to mock expressing cells (**Figure 5**), which suggests Nb9 may be the most promising among the stabilizing one in live cells. Of interest, no effects on GCase activity was observed when lysosome-targeted Nb1, Nb4 and Nb9 were expressed (**Figure 4 and 6**). This appears coherent with the fact that these 3 Nbs belong to the GCase stabilizing Class I Nbs and to a lesser extent increase GCase activity *in vitro*. Hence, a possible interpretation is that when specifically targeted to the lysosomes the primary stabilizing function of the class I Nbs is irrelevant in these organelles. Our model, implying that the Class I Nbs (Nb1, Nb4, Nb9) aid in the trafficking of GCase to the lysosomes, is also in good agreement with the GCase-Nb1 structure, which shows that these Nbs would not interfere with LIMP2 and Saposin C binding.

Additionally, nearly all Nbs that bind wild-type GCase also show binding to the major disease-linked N370S mutant in ELISA and BLI (**Figure 1** and **Supplementary Figure S9)**. This is an important finding with respect to any future therapeutic applications in a clinically relevant setting. Remarkably, Nb10 even displays a >5-fold higher affinity for the N370S mutant compared to wild type GCase. Although, more research is required to make strong conclusions, this finding suggests that Nb10 specifically recognizes a GCase conformation that is more prevalent in the N370S mutant. Interestingly, when measuring *in vitro* N370S GCase activity Nb10 and Nb16 show a significant increase in both the experimental conditions tested (**Figure 6**), further supporting the idea that Nb10 may be more relevant to this mutant than to the wild type enzyme. We also showed that the lysosomal GCase activity of the N370S mutant, when expressed in cells overexpressing the stabilizing Nb4 and Nb9 targeted to the ER, can be increased by about 60-70%. This is coherent with the fact that N370S trafficking was shown to be impaired^13^, suggesting that the mechanism by which ER-Nb4 and ER-Nb9 impact on the mutant enzyme is by promoting its localization at the lysosomes, as we showed for the wild type protein (**Figure 5**). This is enough to increase the lysosomal N370S GCase activity and possibly may be able to reduce ER stress, which will need to be verified by further studies.

In conclusion, upon comparing the overall *in vitro* and *in cellulo* behaviour of all GCase-Nbs, taking into account properties of binding and effects on activity and stability, Nb1, Nb4, and Nb9 emerge as the most promising candidates, with the most remarkable effects seen for the Nb9, both for the wild type and the N370S GCase mutant. All three Nbs bind to the same allosteric binding site, suggesting a crucial role of this site in protein stability and activity. This aligns also with the observation that the most desirable sub-cellular targeting for these Nbs is the ER. This targeting strategy has the potential to counteract the accumulation of unfolded or mutated GCase in the ER, thereby mitigating subsequent ER stress, and concurrently improving the enzyme’s trafficking to the lysosomes, where its activity can be enhanced. Taken together, our results present these Nbs and the pocket they are targeting as valuable starting points for the development of new generations of allosteric molecular chaperones in the treatment of GCase-associated GD and PD.

## Data availability

Atomic coordinates and structure factors for the reported crystal structure have been deposited with the Protein Data Bank under accession code 9ENA.

## Acknowledgments

This work was supported by the Michael J. Fox Foundation for Parkinson’s Research (grant numbers MJFF-17240 and MJFF-020706), the Fonds voor Wetenschappelijk Onderzoek (G031324N to W.V), and a Strategic Research Program Financing from the VUB (SRP50 and SRP95 to W.V. and S.B.). We acknowledge Instruct-ERIC and the FWO for their support to the Nanobody discovery.

We are very grateful to Dr. Prof. Ari Zimran for providing the recombinant commercial enzyme Velaglucerase and Cerezyme for llama immunization and biochemistry experiments. We would like to thank Prof. Friederike Zunke for kindly providing the plasmids for the expression of GCase mutants. We would like to thank Siemen Claeys for excellent technical support, and all members of the Versées lab for comments and discussions.

## Author contributions

N.P. and W.V. conceived the work, contributed to the interpretation and the design of the experiments and wrote the manuscript. T.D.M. performed the nanobody discovery experiments, the biophysical assays, solved the structure, analysed the data and aided in writing the manuscript. C.S. performed the in cell work, analysed the data and aided in writing the manuscript. E.G. adviced on the plasmid generation for the in cell work, contributed to the discussion and wrote the manuscript. J.S. and E.P. supervised the immunizations and adviced on nanobody discovery. S.B. contributed to conveiving the study. G.Z. performed the in vitro activity experiments and the purification of the N370S mutant. I.T. designed and prepared the vector for the transfection of the Nbs in mammalian cells.

## Material and methods

### Immunization and Nb selection

A llama was immunized using a six-week protocol with weekly immunizations of GCase (Velaglucerase, VRPIV®, Takeda) in the presence of GERBU adjuvant, and blood was collected 4 days after the last injection. All animal vaccinations were performed in strict accordance with good practices and EU animal welfare legislation. The construction of immune libraries and Nb selection via phage display were performed using previously described protocols^66^. In brief, starting from the PBMCs collected from the llama blood after immunization, the open reading frames coding for the variable domains of the heavy-chain antibody repertoire were cloned in a pMESy4 phagemid vector (GenBank KF415192), resulting in an immune library of 3.3×10^8^ transformants. This Nb repertoire was expressed on the tip of filamentous phages after rescue with the VCSM13 helper phage. Four phage display selections (two rounds each) were performed using solid phase coating on a 96-well MaxiSorp NUNC-Immuno plate (Thermo Fischer Scientific): (1) glycosylated GCase, (2) glycosylated GCase bound to CBE, (3) deglycosylated GCase, (4) deglycosylated GCase bound to CBE. For selections (2) and (4), proteins were incubated for 30 min on ice with 10 µM conduritol-β-epoxide (CBE). Solid phase coating of all proteins was performed in a coating buffer containing 100 mM NaHCO3 pH8.2. Washing steps were performed with McIlvaine buffer (100 mM Na2HPO4 and 10 mM citric acid) pH5.4, while 0.4% milk was added to this buffer for the binding step. Several single colonies were picked after each round of phage display selection and sequence analysis was used to classify the resulting Nb clones in sequence families based on their CDR3 sequence.

### Nb cloning, expression and purification

After Nb selection, the Nb-coding open reading frames were recloned from the pMESy4 to the pHEN29 vector^66^. Upon expression in an *E. coli* non-suppressor strain both vectors yield proteins with an N-terminal pelB signal sequence to translocate the recombinant protein to the periplasm. Additionally, pMESy4 provides a C-terminal His6-tag and EPEA-tag (= CaptureSelect™ C-tag), while pHEN29 provides a C-terminal LPETGG-His6-EPEA-tag that allows site-specific labelling of the proteins using Sortase chemistry^67^. Expression of Nbs from the former plasmid is used for subsequent structural biology purposes, while the latter is used for labelling of Nbs in BLI experiments.

The Nbs were expressed and purified as previously described^38^. The Nb expression plasmids were transformed in *E. coli* WK6 (Su^-^) cells. Cells were grown at 37°C in Terrific Broth medium to an OD600 ∼1.0. Protein expression was induced by adding 1 mM IPTG (isopropyl β-D-1-thiogalactopyranoside) and incubated overnight at 28°C. After harvesting, cells were lysed by osmotic shock to recover the periplasmic fraction. Nbs are purified via affinity chromatography on Ni^2+^-NTA Sepharose followed by a dialysis step (50 mM HEPES pH 8.0 and 300 mM NaCl).

### Deglycosylation of GCase

Velaglucerase (VRPIV®, Takeda) and Imiglucerase (Cerezyme®, Sanofi) were deglycosylated using PNGase F enzyme (New England Biolabs). The deglycosylation reaction was performed according to the recommendation from the manufacturer, using 0.5 U PNGase F / 1 µg glycosylated protein for 72 hours at 25°C. The deglycosylation reaction was confirmed by SDS-PAGE, and deglycosylated proteins were further purified using size exclusion chromatography (S75 10300 increase GL) in 10 mM MES pH6.5, 100 mM NaCl, 1 mM DTT and 5% glycerol.

### Enzyme-linked immunosorbent assays (ELISA)

GCase, deglycosylated GCase and the GCase N370S mutant were solid-phase coated on the bottom of a 96-well ELISA plates (Maxisorp Nunc-immuno plate, Thermo Fischer Scientific), using a concentration of 1 µg/mL protein in coating buffer (100 mM NaHCO3 pH8.2). All the washing, binding (0.4% milk) and blocking (4% milk) steps were performed using PBS supplemented with 0.05% tween-20. The binding of the Nbs to the coated protein was detected via their EPEA-tag using a 1:4000 CaptureSelect^TM^ Biotin anti-C-tag conjugate (Thermo Fischer Scientific) in combination with 1:1000 Streptavidin Alkaline Phosphatase (Promega). Colour was developed by adding 100 µL of 3 mg/mL disodium 4-nitrophenyl phosphate solution (DNPP, Sigma Aldrich) dissolved in 100 mM Tris, 100 mM NaCl and 5 mM MgCl2 pH9.5, and measured at 405 nm.

### Biolayer Interferometry (BLI)

Prior to BLI experiments, Nbs expressed and purified from the pHEN29 plasmid (with a C-terminal LPETGG-His6-EPEA tag) were site-specifically labeled at their C-terminus using Sortase-mediated exchange with a biotin-labeled GGGYK peptide (GenicBio). GCase was randomly labeled on lysine residues using the EZ-Link™ Sulfo-NHS-LC-Biotin kit (Thermo Fischer Scientific), according to the manufacturers’ recommendations. BLI measurements were performed using an Octet Red96 (FortéBio, Inc.) system in PBS pH7.5, 0.01% Tween-20 supplemented with 0.1% BSA. Either biotinylated GCase or biotinylated Nbs were loaded onto streptavidin-coated (SA) biosensors at a concentration of 1 µg/mL, and the binding of a concentration gradient of unlabelled Nbs or GCase, respectively, was assessed. The association/dissociation traces were fitted with a 1:1 binding model using either the local, partial or global (full) options (implemented in the FortéBio Analysis Software). The resulting Req values were subsequently plotted against the Nb concentration and used to derive the KD values from the corresponding dose-response curves fitted on a Langmuir model (figures were generated using GraphPad Prism10).

### Thermal Shift assays (TSA)

Thermal unfolding of GCase was followed by thermal shift assays using SYPRO orange fluorescence as dye and a CFX connect real-time PCR system (Bio-Rad)^48^. 0.25 mg/mL (4µM) GCase was incubated with 20 µM of the different Nbs for 30 min on ice and combined with 5x SYPRO Orange protein Gel Stain (Thermo Fischer Scientific), in Mc Ilvaine buffer pH 5.2 or pH 7.0 consisting of 0.1 M citric acid and 0.2 M dibasic sodium phosphate in a total volume of 25 µL. The temperature was increased from 20°C to 80°C at 1°C/min steps. All measurements were performed in triplicate and the melting temperatures were determined by fitting the first derivatives of the data with a Boltzmann sigmoidal equation (GraphPad Prism). ΔTm (°C) values are the difference in melting temperature between GCase and GCase incubated with different Nbs.

### Structure determination and analysis

Imiglucerase (Cerezyme®) was partially deglycosylated prior to crystallization as previously described^50^. Crystals of this protein in complex with Nb1 were obtained by co-crystallization using the sitting-drop vapor diffusion method at 277K. GCase (Imiglucerase) and Nb1 proteins were mixed to obtain a concentration of 10 mg/mL and 3.5 mg/mL respectively (1:1.2 molar ratio). Crystals were obtained in a crystallization solution of 1.6 M Magnesium sulfate heptahydrate, 0.1 M MES pH6.5. Crystals were cryo-protected in mother liquor supplemented with 25% glycerol.

Data were collected at 100K at the Proxima 2 beamline of the Soleil synchrotron (λ = 0.9801 Å), to a resolution of 1.7 Å. Diffraction data were integrated and scaled with autoPROC^68^, using the default pipeline including XDS, Truncate, Aimless and STARANISO. Crystal belonged to the tetragonal space group I422. The structure was solved using the molecular replacement method based on PDB 2J25 for GCase and 7A17 (chain B) for Nb1, and refined with the phenix.refine module (Phenix version 1.20.1)^69^ alternated with manual building in Coot^70^. **Supplementary Table S1** summarizes data collection, processing and refinement statistics. Structure and structure factors were deposited in the PDB (code 9ENA). All structural figures were produced with PyMOL (The PyMOL Molecular Graphics System, Version 2.3.3, Schrödinger, LLC.).

### Cell culture, plasmids and transfection

Human embryonic kidney cells (HEK293T) and human fibroblast cells were cultured in Dulbecco’s modified Eagle’s medium (DMEM, Sigma Aldrich), supplemented with fetal bovine serum (FBS, 10% Sigma Aldrich) and 100 U/mL penicillin, and 100 μg/mL streptomycin (Sigma Aldrich) at 37°C and 5% CO2. For transient transfection, polyethyleneimine (PEI) was used as a transfection reagent and HEK293T cells were incubated with DNA:PEI (1:2) in OPTIMEM (Life Technologies) for two hours, before media change. After 48 h, activity assays, western blotting analyses or imaging experiments were performed.

### Mice colony

*Gba1*^-/-^ *h*N370S mice were purchased from the Jackson laboratories (https://www.jax.org/strain/032791). Mouse genotyping was performed with WONDER Taq Hot START (Euroclone) using the following primers: 5’-TCCTCACCTCCTCAGATGCT-3’ (mutant forward), 5’-ACCCTCGGGTTTTAAGCTG-3’ (mutant reverse), 5’-CTCTGCAGTTGTGGTCGTGT-3’ (wild-type forward), 5’-GTCCATGCTAAGCCCAGGT-3’ (wild-type reverse), 5’-CTG TCC CTG TAT GCC TCT GG-3’ (Internal Positive Control Forward), 5’-AGATGGAGAAAGGACTAGGCTACA-3’ (Internal Positive Control Reverse), 5’-CAG CCA TGA TGC TTA CCC TAC-3’ (Transgene Reverse), 5’-GCT AAC CAT GTT CAT GCC TTC-3’ (Transgene forward).

Animals were maintained and experiments were conducted according to the Italian Ministry of Health and the approval by the Ethical Committee of the University of Padova (authorization number D2784.N.QHV).

### Measurement of GCase activity in live cells

A selective lysosomal GCase substrate, 5-(Pentafluorobenzoylamino) Fluorescein Di-β-D-Glucopyranoside (PFB-FDGlu, ThermoFisher Scientific) was used to evaluate GCase activity in live cells. HEK293T cells were cultured in a 24-well plate (150000 cells/well) and transfected with mCherry plasmids (1 µg DNA/well). After 48 h, PFB-FDGlu (50 µg/mL) was added for 6 h and the nPFB-FDGlu fluorescence was measured by BD FACSAria™ III Cell Sorter (λex 492 nm and λem 516 nm).

Gcase activity was also evaluated in *GBA1* knockdown HEK293T (11F clone) overexpressing the N370S GCase mutant. Briefly, the cells were cultured in 24-well plate (150000 cells/well) and were co-transfected with mCherry plasmids (Mock, Nb1, Nb4 and Nb9) and pCMV3_ N370S_His plasmid encoding the N370S GCase mutant (1 µg DNA/well).

After 48 h, PFB-FDGlu (50 µg/mL) was added for 6 h and the nPFB-FDGlu fluorescence was measured by BD FACSAria™ III Cell Sorter (λex 492 nm and λem 516 nm). Non-transfected wild type and *Gba1* knockdown cells were used as controls.

### Measurement of GCase activity in cell or tissue lysates

GCase activity was measured in HEK293T transfected with different ER- or lysosomal targeting Nbs by using the 4-methylumbelliferyl-β-D-glucopyranoside (4-MU) assay. Cells were cultured in a 12-well plate (300000 cells/well) and transfected with eGFP plasmids (2 DNA µg/well). After 48h, cells were lysed in RIPA buffer supplemented with protease inhibitors (Roche) and protein was quantified by Pierce® Bicinchoninic acid (BCA) Protein Assay Kit assay.

As for tissues, they were isolated from 8-month-old *Gba1*^-/-^ *h*N370S transgenic mice and stored at −80°C. Then, tissues were lysed in RIPA buffer (ratio 1:4) supplemented with protease inhibitors (Roche) and protein was quantified by BCA Protein Assay Kit assay.

Cell lysate samples were prepared in citrate phosphate buffer pH 4.5 (0.1 M Citric Acid, 0.2 M Na2HPO4) (20 µL, 2 µg of protein) and incubated with the 4-MU substrate (3 mM in citrate phosphate buffer with 0.2% taurocholate) for 90 min at 37°C. Tissue lysates were prepared in the same manner but they were preincubated with Nbs (2.5 µM) for 30’ at 37 °C before starting the incubation with the substrate. The reaction was stopped by adding 240 µL of stop buffer (0.2 M NaOH, 0.2 M Glycine pH 10). Fluorescence was measured in a fluorescence microplate reader (Victor X3, Perkin Elmer). The assay was performed in triplicate.

### Western blot analysis

HEK293T cells were cultured in a 12-well plate (300000 cells/well) and transfected with eGFP plasmids (2 µg DNA/well). After 48h, cells were lysed in RIPA buffer supplemented with protease inhibitors cocktail (Roche). Lysates were centrifuged at 20000 × g at 4°C. Protein concentration was determined using the Pierce® BCA Protein Assay Kit. Equal amounts of protein were loaded on gradient gel 4-20%. Tris-MOPS-SDS gels (GenScript). The resolved proteins were then transferred to PVDF membranes (BioRad), through a semi-dry Trans Blot® TurboTM Transfer System (BioRad). PVDF membranes were blocked in Tris-buffered saline plus 0.1% Tween (TBS-T) and 5% non-fat dry milk for 1 h and then incubated overnight at 4°C with primary antibodies diluted in TBS-T plus 5% non-fat milk. The following primary antibodies were used: mouse anti-β-actin (1:10000, A2066 Sigma-Aldrich), rabbit anti-calnexin (1:1000, ab22595 Abcam), mouse anti-LAMP1(1:400, sc-20011 Santa Cruz Biotechnology), anti-FLAG HRP (1:1000, A8592 Sigma-Aldrich), rabbit anti-GBA1 (1:1000, G4171 Sigma-Aldrich). After incubation with horse-radish peroxidase (HRP)-conjugated secondary antibodies (goat anti-rabbit-HRP and goat anti-mouse-HRP, Sigma-Aldrich) at room temperature for 1 h, immunoreactive proteins were visualized using Immobilon® Classico Western HRP Substrate (Millipore) or Immobilon® Forte Western HRP Substrate (Millipore) by Imager CHEMI Premium detector (VWR). The densitometric analysis of the detected bands was performed by using the IMAGE J software.

### Lysosomal proteolytic activity

HEK293T cells were plated in a 24-well plate and transfected with the selected ER- or lysosomal targeting Nbs (1 µg/well). Lysosomal protease activity was evaluated by using DQ-Red BSA dye, a fluorogenic substrate for proteases that is hydrolyzed in acidic, hydrolase-active endo-lysosomes to smaller protein fluorescent peptides. Cells were incubated with DQ-Red BSA (10 µg/mL) for 30 minutes. Then, fresh medium was added and fluorescence was measured after 2h by BD LSR Fortessa™ X-20 Cell Analyzer (λex 590 nm and λem 620 nm). Chloroquine (CQ 50 µM), a lysosomotropic agent that blocks endosomal acidification, was used as a positive control.

### ENDO H and PNGase F treatment of cell lysates

HEK293T cells were transfected with ER-targeting Nb1, Nb4 and Nb9 (500000 cells/well, 3 µg DNA/well) and then lysed in RIPA buffer, as described above. 30 µg of proteins from lysed cells were digested with 500 units of endoglycosidase H (ENDO H, Promega) and 10 units of Peptide N-glycosidase F (PNGase F, Promega) enzymes according to manufacturer’s instructions, after which they were used for immunoblotting. Non-transfected cells subjected to all digestion steps without enzymes were used as a positive control.

### Immunofluorescence image analysis

HEK293T cells were cultured in 24-well (50000 cells/well) and transfected with ER- and lysosomal targeting nanobodies (1 µg DNA/well). Cells were fixed with 4% paraformaldehyde (PFA, Sigma-Aldrich) for 20 min, permeabilized with PBS-0.1% TritonTM X-100 for 20 minutes and blocked with PBS-5% fetal bovine serum (FBS) for 1 h. Primary and secondary antibodies were prepared in a blocking solution (1:200 in PBS-5% FBS). The following antibodies were used: anti-FLAG antibody (F7425, Sigma-Aldrich) together with the anti-calnexin (ab22595, Sigma-Aldrich) or the anti-LAMP2A (ab18528, Abcam), Goat anti-Rabbit Alexa Fluor 633 (A21071, ThermoFisher Scientific) and Goat anti-Mouse Alexa Fluor™ 568 (A11004, ThermoFisher Scientific).

Nuclei were stained with Hoechst 33258 pentahydrate (bis-benzimide) (Invitrogen, 1:10000 in H2O). The images were acquired using a 63x magnification objective by means of a Zeiss LSM700 laser scanning confocal microscope. The co-localization of ER-targeting Nbs in the ER of transfected cells was evaluated by calculating the Pearson coefficient using a colocalization plugin (Jacop) of Image J software.

### N370S GCase purification

Recombinant GCase N370S mutant was produced by using 293T Freestyle cells (Thermo Fisher Cat. No. R79007) grown in suspension in 293 Freestyle medium (Thermo Fisher). Cells were grown up to 10^6^ cells/ml at 37°C CO2 8% in agitation at 125 rpm, then transfected using 1 µg of pCMV_GBA_N370S_His and 3µg of PEI per 10^6^ cells in OPTIMEM (Life Technologies). 12 hours after transfection, 3.5 mM valproic acid dissolved in water was added to the cell suspension and incubation was continued for 96 hours. Then, the medium containing the secreted N370S GCase protein was collected and clarified by centrifugation at 500 x g for 20 min at 4°C. The collected supernatant was filtered at 4°C with a 0.22 µm filter. The medium was then loaded on a His-select Nickel affinity column (Millipore) equilibrated in 50 mM NaH2PO4, 300 mM NaCl, 10 mM imidazole pH 8. The column was then washed with 50 mM NaH2PO4, 300 mM NaCl, 30 mM imidazole pH 8 and the protein was eluted with 50 mM NaH2PO4, 300 mM NaCl, 30 mM imidazole pH 8. Buffer exchange was performed via PD-10 desalting column (GE Healthcare) to 50mM MES pH 5.0 for experiments and storage. Protein quality was confirmed by SDS-PAGE and western blot analysis.

### Measurement of the activity of recombinant GCase

GCase activity was measured on the commercially available enzyme (Velaglucerase) and on recombinant N370S GCase. Briefly, protein samples were dissolved in citrate phosphate buffer pH 4.5 (0.1 M Citric Acid, 0.2 M Na2HPO4) (20 µL, 2 µg of protein) and reincubated with Nbs (2.5 µM) for 30’ at 37 °C. Incubation with the 4-MU substrate (3 mM in citrate phosphate buffer with 0.2% taurodeoxycholate, Sigma-Aldrich) for 90 min at 37°C. The reaction was stopped by adding 240 µL of stop buffer (0.2 M NaOH, 0.2 M Glycine pH 10). Fluorescence was measured in a fluorescence microplate reader (Victor X3, Perkin Elmer). The assay was performed in triplicate or quadruplicate.

### Statistical analysis

Independent experiments and technical replicates were reported in the figure legends. Statistical analysis and graphical visualization were carried out with GraphPad Prism Software Inc. (Version 8) and details about the analysis are also reported in the figure legends.

### Nbs cloning in vectors for the expression in mammalian cells

For the expression of Nbs in mammalian cells, two different plasmid backbones were used, pRP[Exp]-Hygro-CMV for transfection and pLV[Exp]-Puro-EF1A for lentivirus production; both were customized and purchased from VectorBuilder Inc.

Plasmids were designed for each Nb to be expressed with a 3xFLAG epitope at the C-terminus and a sequence targeting the Endoplasmic Reticulum (ER) or the Lysosome (LYSO). Furthermore, the expression via IRES of a fluorescent protein (EGFP or mCherry) was included in order to easily identify transfected/transduced cells. VectorBuilder provided plasmids with a Stuffer ORF sequence (peptide of E. coli beta-galactosidase) flanked by NheI and BstBI restriction sites.

The ER targeting sequence was chosen among those of eukaryotic proteins that are known to target the ER and whose transport mechanisms are known. In particular, MKWVTFLLLLFISAFSR is the Preproalbumin signal peptide^55^ and was inserted upstream of the NheI site in the ER plasmid series.

The targeting of proteins to the lysosome has a more complex mechanism and we lack a single and well-defined signal sequence. In the work of Fan X. et al, 2014^54^ the authors demonstrated the possibility of targeting a protein of interest to the lysosomes by fusing it at the C-terminus with a sequence obtained from the sum of 3 different signal sequences known for Chaperone-mediated autophagy (CMA-targeting motif, CTM). Therefore, we used the same sequence, KFERQKILDQRFFE, as a signal peptide for the lysosome. The sequence was included in frame following 3XFLAG in the LYSO plasmid series.

All Nbs sequences of interest were amplified by PCR with the same primers, designed on the conserved regions at the N-term and C-term and including the NheI and BstBI restriction sites, respectively. The plasmids were all digested with these endonucleases to eliminate the Stuffer ORF and ligated with the sequences of interest. The original plasmids with Stuffer ORF were used as controls for the effect of the transfection.

List of plasmids:

pRP[Exp]-Hygro-CMV>{ER-ORF_Stuffer/3xFLAG}:IRES:EGFP (VectorBuilder Inc.)
pRP[Exp]-Hygro-CMV>{ER-ORF_Stuffer/3xFLAG}:IRES:mCherry (VectorBuilder Inc.)
pRP[Exp]-Hygro-CMV>{ORF_Stuffer/3xFLAG-LYSO}:IRES:EGFP (VectorBuilder Inc.)
pRP[Exp]-Hygro-CMV>{ORF_Stuffer/3xFLAG-LYSO}:IRES:mCherry (VectorBuilder Inc.)
pRP[Exp]-Hygro-CMV>{ER-ORF_NbXX/3xFLAG}:IRES:EGFP
pRP[Exp]-Hygro-CMV>{ER-ORF_NbXX/3xFLAG}:IRES:mCherry
pRP[Exp]-Hygro-CMV>{ORF_NbXX /3xFLAG-LYSO}:IRES:EGFP
pRP[Exp]-Hygro-CMV>{ORF_NbXX/3xFLAG-LYSO}:IRES:mCherry
pLV[Exp]-Puro-EF1A>{ER-ORF_Stuffer/3xFLAG}:IRES:mCherry (VectorBuilder Inc.)
pLV[Exp]-Puro-EF1A>{ORF_Stuffer/3xFLAG-LYSO}:IRES:mCherry (VectorBuilder Inc.)
pLV[Exp]-Puro-EF1A>{ER-ORF_NbXX/3xFLAG}:IRES:mCherry
pLV[Exp]-Puro-EF1A>{ORF_NbXX/3xFLAG-LYSO}:IRES:mCherry

### *GBA1* knockdown in HEK293T cells via Crispr/Cas9 and stable cell lines generation

GBA1 knockdown in HEK293T cells via Crispr/Cas9 and stable cell lines generation The CRISPR/Cas9 system was used to silence the *GBA1* gene in HEK293T cells. Two single guide RNA (sgRNA) sequences were selected. The sgRNA sequences targeting *GBA1* were:

GBA1_sgRNA(1)_FOR: CACCGATGATGCTTACCCTACTCAA
GBA1_sgRNA(1)_REV: aaacTTGAGTAGGGTAAGCATCATC
GBA1_sgRNA(2)_FOR: CACCGCGCTATGAGAGTACACGCAG
GBA1_sgRNA(2)_REV: aaacCTGCGTGTACTCTCATAGCGC

To establish a *Gba1* knockdown clone, HEK293T cells were seeded in 6-well plate (450000 cells/well). Then, cells were transfected wih two different plasmids (guide 1 and guide 2) (4µg DNA/well, DNA:PEI 1:2 in opti-MEM). Antibiotic selection (puromycin 2.5 µg/mL) was started after 24h and continued for at least three days. Different single clones were selected, expanded and then used for biological assays.

## References

1 Marques ARA, Mirzaian M, Akiyama H, Wisse P, Ferraz MJ, Gaspar P et al. Glucosylated cholesterol in mammalian cells and tissues: Formation and degradation by multiple cellular ô -glucosidases. J Lipid Res 2016; 57: 451–463.

2 Grabowski GA. Phenotype, diagnosis, and treatment of Gaucher’s disease. Lancet 2008; 372: 1263– 1271.

3 Sidransky E, Nalls MA, Aasly JO, Aharon-Peretz J, Annesi G, Barbosa ER et al. Multicenter Analysis of Glucocerebrosidase Mutations in Parkinson’s Disease. N Engl J Med 2009; 361: 1651–1661.

4 Daykin EC, Ryan E, Sidransky E. Diagnosing neuronopathic Gaucher disease: New considerations and challenges in assigning Gaucher phenotypes. Mol Genet Metab 2021; 132: 49–58.

5 Roshan Lal T, Sidransky E. The Spectrum of Neurological Manifestations Associated with Gaucher Disease. Diseases 2017; 5: 10.

6 Furderer ML, Hertz E, Lopez GJ, Sidransky E. Neuropathological Features of Gaucher Disease and Gaucher Disease with Parkinsonism. Int J Mol Sci 2022; 23. doi:10.3390/ijms23105842.

7 Kalia L V., Lang AE. Parkinson’s disease. Lancet 2015; 386: 896–912.

8 Neumann J, Bras J, Deas E, O’sullivan SS, Parkkinen L, Lachmann RH et al. Glucocerebrosidase mutations in clinical and pathologically proven Parkinson’s disease. Brain 2009; 132: 1783–1794.

9 Clark LN, Ross BM, Wang Y, Mejia-Santana H, Harris J, Louis ED et al. Mutations in the glucocerebrosidase gene are associated with early-onset Parkinson disease. Neurology 2007; 69: 1270–1277.

10 Stoker TB, Stoker TB, Camacho M, Winder-Rhodes S, Liu G, Liu G et al. Impact of GBA1 variants on long-term clinical progression and mortality in incident Parkinson’s disease. J Neurol Neurosurg Psychiatry 2020; 91: 695–702.

11 Pokorna S, Khersonsky O, Lipsh-Sokolik R, Goldenzweig A, Nielsen R, Ashani Y et al. Design of a stable human acid-β-glucosidase: towards improved Gaucher disease therapy and mutation classification. FEBS J 2023; 290: 3383–3399.

12 Reczek D, Schwake M, Schröder J, Hughes H, Blanz J, Jin X et al. LIMP-2 Is a Receptor for Lysosomal Mannose-6-Phosphate-Independent Targeting of β-Glucocerebrosidase. Cell 2007; 131: 770–783.

13 Fernandes HJR, Hartfield EM, Christian HC, Emmanoulidou E, Zheng Y, Booth H et al. ER Stress and Autophagic Perturbations Lead to Elevated Extracellular α-Synuclein in GBA-N370S Parkinson’s iPSC-Derived Dopamine Neurons. Stem Cell Reports 2016; 6: 342–356.

14 Malini E, Grossi S, Deganuto M, Rosano C, Parini R, Dominisini S et al. Functional analysis of 11 novel GBA alleles. Eur J Hum Genet 2014; 22: 511–516.

15 Smith L, Mullin S, Schapira AHV. Insights into the structural biology of Gaucher disease. Exp Neurol 2017; 298: 180–190.

16 Osellame LD, Rahim AA, Hargreaves IP, Gegg ME, Richard-Londt A, Brandner S et al. Mitochondria and quality control defects in a mouse model of gaucher disease - Links to parkinson’s disease. Cell Metab 2013; 17: 941–953.

17 Plotegher N, Perocheau D, Ferrazza R, Massaro G, Bhosale G, Zambon F et al. Impaired cellular bioenergetics caused by GBA1 depletion sensitizes neurons to calcium overload. Cell Death Differ 2020; 27: 1588–1603.

18 Gegg ME, Burke D, Heales SJR, Cooper JM, Hardy J, Wood NW et al. Glucocerebrosidase deficiency in substantia nigra of parkinson disease brains. Ann Neurol 2012; 72: 455–463.

19 Gegg ME, Sweet L, Wang BH, Shihabuddin LS, Sardi SP, Schapira AHV. No evidence for substrate accumulation in Parkinson brains with GBA mutations. Mov Disord 2015; 30: 1085–1089.

20 Giladi N, Alcalay RN, Cutter G, Gasser T, Gurevich T, Höglinger GU et al. Safety and efficacy of venglustat in GBA1-associated Parkinson’s disease: an international, multicentre, double-blind, randomised, placebo-controlled, phase 2 trial. Lancet Neurol 2023; 22: 661–671.

21 Steet RA, Chung S, Wustman B, Powe A, Do H, Kornfeld SA. The iminosugar isofagomine increases the activity of N370S mutant acid β-glucosidase in Gaucher fibroblasts by several mechanisms. Proc Natl Acad Sci U S A 2006; 103: 13813–13818.

22 Zheng W, Padia J, Urban DJ, Jadhav A, Goker-Alpan O, Simeonov A et al. Three classes of glucocerebrosidase inhibitors identified by quantitative high-throughput screening are chaperone leads for Gaucher disease. Proc Natl Acad Sci U S A 2007; 104: 13192–13197.

23 Zheng J, Chen L, Skinner OS, Ysselstein D, Remis J, Lansbury P et al. β-Glucocerebrosidase Modulators Promote Dimerization of β-Glucocerebrosidase and Reveal an Allosteric Binding Site. J Am Chem Soc 2018; 140: 5914–5924.

24 Mazzulli JR, Zunke F, Tsunemi T, Toker NJ, Jeon S, Burbulla LF et al. Activation of β-glucocerebrosidase reduces pathological α-synuclein and restores lysosomal function in Parkinson’s patient midbrain neurons. J Neurosci 2016; 36: 7693–7706.

25 Patnaik S, Zheng W, Choi JH, Motabar O, Southall N, Westbroek W et al. Discovery, structure - Activity relationship, and biological evaluation of noninhibitory small molecule chaperones of glucocerebrosidase. J Med Chem 2012; 55: 5734–5748.

26 Migdalska-Richards A, Ko WKD, Li Q, Bezard E, Schapira AHV. Oral ambroxol increases brain glucocerebrosidase activity in a nonhuman primate. Synapse 2017; 71: 17–19.

27 Migdalska-Richards A, Daly L, Bezard E, Schapira AHV. Ambroxol effects in glucocerebrosidase and α-synuclein transgenic mice. Ann Neurol 2016; 80: 766–775.

28 McNeill A, Magalhaes J, Shen C, Chau KY, Hughes D, Mehta A et al. Ambroxol improves lysosomal biochemistry in glucocerebrosidase mutation-linked Parkinson disease cells. Brain 2014; 137: 1481– 1495.

29 Kopytova AE, Rychkov GN, Nikolaev MA, Baydakova G V., Cheblokov AA, Senkevich KA et al. Ambroxol increases glucocerebrosidase (GCase) activity and restores GCase translocation in primary patient-derived macrophages in Gaucher disease and Parkinsonism. Park Relat Disord 2021; 84: 112–121.

30 Pantoom S, Hules L, Schöll C, Petrosyan A, Monticelli M, Pospech J et al. Mechanistic Insight into the Mode of Action of Acid β-Glucosidase Enhancer Ambroxol. Int J Mol Sci 2022; 23. doi:10.3390/ijms23073536.

31 Muyldermans S. Nanobodies: Natural single-domain antibodies. Annu Rev Biochem 2013; 82: 775–797.

32 Jovčevska I, Muyldermans S. The Therapeutic Potential of Nanobodies. BioDrugs 2020; 34: 11–26.

33 De Genst E, Silence K, Decanniere K, Conrath K, Loris R, Kinne J et al. Molecular basis for the preferential cleft recognition by dromedary heavy-chain antibodies. Proc Natl Acad Sci U S A 2006; 103: 4586–4591.

34 Steyaert J, Kobilka BK. Nanobody stabilization of G protein-coupled receptor conformational states. Curr Opin Struct Biol 2011; 21: 567–572.

35 Schubert AF, Gladkova C, Pardon E, Wagstaff JL, Freund SMV, Steyaert J et al. Structure of PINK1 in complex with its substrate ubiquitin. Nature 2017; 552: 51–56.

36 Hmila I, Vaikath NN, Majbour NK, Erskine D, Sudhakaran IP, Gupta V et al. Novel engineered nanobodies specific for N-terminal region of alpha-synuclein recognize Lewy-body pathology and inhibit in-vitro seeded aggregation and toxicity. FEBS J 2022; 289: 4657–4673.

37 Paesmans J, Martin E, Deckers B, Berghmans M, Sethi R, Loeys Y et al. A structure of substrate-bound synaptojanin1 provides new insights in its mechanism and the effect of disease mutations. Elife 2020; 9: 1–27.

38 Singh RK, Soliman A, Guaitoli G, Stormer E, von Zweydorf F, Dal Maso T et al. Nanobodies as allosteric modulators of Parkinson’s disease-associated LRRK2. Pnas 2022; 119: 1–34.

39 Brumshtein B, Salinas P, Peterson B, Chan V, Silman I, Sussman JL et al. Characterization of gene-activated human acid-β-glucosidase: Crystal structure, glycan composition, and internalization into macrophages. Glycobiology 2010; 20: 24–32.

40 Zimran A. Velaglucerase alfa: A new option for gaucher disease treatment. Drugs of Today 2011; 47: 515–529.

41 Tekoah Y, Tzaban S, Kizhner T, Hainrichson M, Gantman A, Golembo M et al. Glycosylation and functionality of recombinant β-glucocerebrosidase from various production systems. Biosci Rep 2013; 33: 771–781.

42 Klein AD, Ferreira NS, Ben-Dor S, Duan J, Hardy J, Cox TM et al. Identification of Modifier Genes in a Mouse Model of Gaucher Disease. Cell Rep 2016; 16: 2546–2553.

43 Kedariti M, Frattini E, Baden P, Cogo S, Civiero L, Ziviani E et al. LRRK2 kinase activity regulates GCase level and enzymatic activity differently depending on cell type in Parkinson’s disease. npj Park Dis 2022; 8. doi:10.1038/s41531-022-00354-3.

44 Sanchez-Martinez A, Beavan M, Gegg ME, Chau KY, Whitworth AJ, Schapira AHV. Parkinson disease-linked GBA mutation effects reversed by molecular chaperones in human cell and fly models. Sci Rep 2016; 6. doi:10.1038/srep31380.

45 Alcalay RN, Levy OA, Waters CC, Fahn S, Ford B, Kuo SH et al. Glucocerebrosidase activity in Parkinson’s disease with and without GBA mutations. Brain 2015; 138: 2648–2658.

46 Maegawa GHB, Tropak MB, Buttner JD, Rigat BA, Fuller M, Pandit D et al. Identification and characterization of ambroxol as an enzyme enhancement agent for Gaucher disease. J Biol Chem 2009; 284: 23502–23516.

47 Tran ML, Génisson Y, Ballereau S, Dehoux C. Second-Generation Pharmacological Chaperones: Beyond Inhibitors. Molecules 2020; 25: 3145.

48 Zhang R, Monsma F. Fluorescence-based thermal shift assays. Curr Opin Drug Discov Devel 2010; 13: 389–402.

49 Brumshtein B, Wormald MR, Silman I, Futerman AH, Sussman JL. Structural comparison of differently glycosylated forms of acid-β-glucosidase, the defective enzyme in Gaucher disease. Acta Crystallogr Sect D Biol Crystallogr 2006; 62: 1458–1465.

50 Dvir H, Harel M, McCarthy AA, Toker L, Silman I, Futerman AH et al. X-ray structure of human acid-β-glucosidase, the defective enzyme in Gaucher disease. EMBO Rep 2003; 4: 704–709.

51 Lieberman RL, Wustman BA, Huertas P, Powe AC, Pine CW, Khanna R et al. Structure of acid β-glucosidase with pharmacological chaperone provides insight into Gaucher disease. Nat Chem Biol 2007; 3: 101–107.

52 Lieberman RL. A guided tour of the structural biology of gaucher disease: Acid-β-glucosidase and saposin C. Enzyme Res 2011; 2011. doi:10.4061/2011/973231.

53 Romero R, Ramanathan A, Yuen T, Bhowmik D, Mathew M, Munshi LB et al. Mechanism of glucocerebrosidase activation and dysfunction in Gaucher disease unraveled by molecular dynamics and deep learning. Proc Natl Acad Sci U S A 2019; 116: 5086–5095.

54 Fan X, Jin WY, Lu J, Wang J, Wang YT. Rapid and reversible knockdown of endogenous proteins by peptide-directed lysosomal degradation. Nat Neurosci 2014; 17: 471–480.

55 Kurzchalia T V., Wiedmann M, Girshovicht AS, Bochkarevat ES, Bielka H, Rapoport TA. The signal sequence of nascent preprolactin interacts with the 54K polypeptide of the signal recognition particle. Nature 1986; 320: 634–636.

56 Singh B, Bhaskar S. Methods for Detection of Autophagy in Mammalian Cells. In: Stem Cells and Aging. Methods in Molecular Biology, vol 2045. 2018.

57 Navarro-Romero A, Fernandez-Gonzalez I, Riera J, Montpeyo M, Albert-Bayo M, Lopez-Royo T et al. Lysosomal lipid alterations caused by glucocerebrosidase deficiency promote lysosomal dysfunction, chaperone-mediated-autophagy deficiency, and alpha-synuclein pathology. npj Park Dis 2022; 8. doi:10.1038/s41531-022-00397-6.

58 Wei RR, Hughes H, Boucher S, Bird JJ, Guziewicz N, Van Patten SM et al. X-ray and biochemical analysis of N370S mutant human acid β-glucosidase. J Biol Chem 2011; 286: 299–308.

59 García-Sanz P, Orgaz L, Bueno-Gil G, Espadas I, Rodríguez-Traver E, Kulisevsky J et al. N370S-GBA1 mutation causes lysosomal cholesterol accumulation in Parkinson’s disease. Mov Disord 2017; 32: 1409–1422.

60 Sanders A, Hemmelgarn H, Melrose HL, Hein L, Fuller M, Clarke LA. Transgenic mice expressing human glucocerebrosidase variants: Utility for the study of Gaucher disease. *Blood Cells*, Mol Dis 2013; 51: 109–115.

61 Magalhaes J, Gegg ME, Migdalska-Richards A, Doherty MK, Whitfield PD, Schapira AHV. Autophagic lysosome reformation dysfunction in glucocerebrosidase deficient cells: Relevance to Parkinson disease. Hum Mol Genet 2015; 25: 3432–3445.

62 Kinghorn KJ, Grönke S, Castillo-Quan JI, Woodling NS, Li L, Sirka E et al. A Drosophila model of neuronopathic gaucher disease demonstrates lysosomal-autophagic defects and altered mTOR signalling and is functionally rescued by rapamycin. J Neurosci 2016; 36: 11654–11670.

63 Mazzulli JR, Xu YH, Sun Y, Knight AL, McLean PJ, Caldwell GA et al. Gaucher disease glucocerebrosidase and α-synuclein form a bidirectional pathogenic loop in synucleinopathies. Cell 2011; 146: 37–52.

64 Martínez-Bailén M, Clemente F, Matassini C, Cardona F. GCase Enhancers: A Potential Therapeutic Option for Gaucher Disease and Other Neurological Disorders. Pharmaceuticals 2022; 15. doi:10.3390/ph15070823.

65 Zunke F, Andresen L, Wesseler S, Groth J, Arnold P, Rothaug M et al. Characterization of the complex formed by β-glucocerebrosidase and the lysosomal integral membrane protein type-2. Proc Natl Acad Sci 2016; 113: 3791–3796.

66 Pardon E, Laeremans T, Triest S, Rasmussen SGF, Wohlkönig A, Ruf A et al. A general protocol for the generation of Nanobodies for structural biology. Nat Protoc 2014; 9: 674–693.

67 Massa S, Vikani N, Betti C, Ballet S, Vanderhaegen S, Steyaert J et al. Sortase A-mediated site-specific labeling of camelid single-domain antibody-fragments: a versatile strategy for multiple molecular imaging modalities. Contrast Media Mol Imaging 2016; 11: 328–339.

68 Vonrhein C, Flensburg C, Keller P, Sharff A, Smart O, Paciorek W et al. Data processing and analysis with the autoPROC toolbox. Acta Crystallogr Sect D Biol Crystallogr 2011; 67: 293–302.

69 Afonine P V., Grosse-Kunstleve RW, Echols N, Headd JJ, Moriarty NW, Mustyakimov M et al. Towards automated crystallographic structure refinement with phenix.refine. Acta Crystallogr Sect D Biol Crystallogr 2012; 68: 352–367.

70 Emsley P, Lohkamp B, Scott WG, Cowtan K. Features and development of Coot. Acta Crystallogr Sect D Biol Crystallogr 2010; 66: 486–501.

