## Supplementary figures for "Identification and characterization of nanobodies acting as molecular chaperones for glucocerebrosidase through a novel allosteric mechanism"

### Supplemental information

**Supplementary Table S1: X-ray data collection and refinement statistics.**

| | |
|---|---|
| PDB code | 9ENA |
| **Data collection** | |
| Synchrotron | Soleil |
| Beamline | Px2A |
| Wavelength (Å) | 0.9801 |
| Resolution range (Å)* | 46.86 - 1.7 (1.76 - 1.7) |
| Space group | I422 |
| Unit cell dimensions (Å) | a = 113.209 |
| | b = 113.209 |
| | c = 257.373 |
| Unit cell angles (º) | α = 90 |
| | β = 90 |
| | γ =90 |
| Spherical completeness (%)* | 85.3 (33.2) |
| Ellipsoidal completeness (%)* | 95.3 (79.9) |
| Unique reflections | 78277 |
| Mean (I)/SD(I)* | 13.4 (1.7) |
| CC (1/2)* | 0.998 (0.520) |
| Multiplicity* | 17.1 (9.9) |
| $R_{meas}$ (%)* | 21.0 (169.8) |
| **Refinement** | |
| Resolution range (Å) | 46.86 - 1.7 (1.761 - 1.7) |
| $R_{work}$ (%) | 16.30 |
| $R_{free}$ (%)† | 19.46 |
| Model content | |
| Molecules per AU | 2 |
| Protein atoms per AU | 4961 |
| Ligand atoms per AU | 84 |
| Metal atoms per AU | 2 |
| Water molecules per AU | 808 |
| Wilson B factors (Å$^2$) | 15.88 |
| Average B factors (Å$^2$) | 20.88 |
| Protein atoms | 18.85 |
| Ligand atoms | 40.79 |
| Metal atoms | 22.49 |
| Water molecules | 31.25 |
| Rmsd bonds (Å) | 0.008 |
| Rmsd angles (º) | 0.96 |
| Ramachandran plot (%) (favored, outliers) | 96.95, 0.48 |

* Values in parentheses are for the high-resolution shell.
† $R_{free}$ is based on a subset of 5% of reflections omitted during refinement.
AU, asymmetric unit.

A

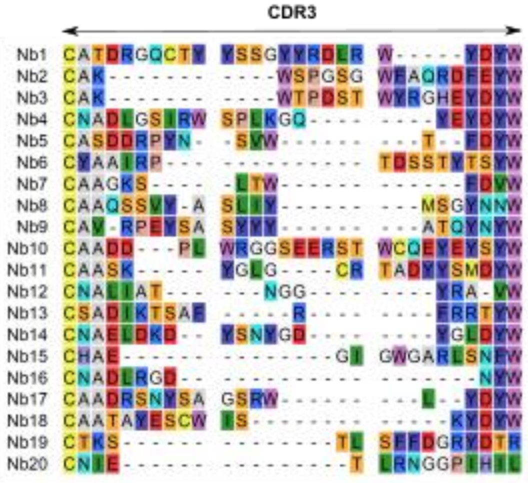


B

| Nb# | Internal database# | Selection |
|---|---|---|
| 1 | CA16655 | deglycosylated GCase bound to CBE |
| 2 | CA16657 | deglycosylated GCase |
| 3 | CA16662 | glycosylated GCase bound to CBE |
| 4 | CA16666 | deglycosylated GCase |
| 5 | CA16668 | deglycosylated GCase bound to CBE |
| 6 | CA16669 | glycosylated GCase |
| 7 | CA16670 | deglycosylated GCase bound to CBE |
| 8 | CA16671 | glycosylated GCase bound to CBE |
| 9 | CA16672 | deglycosylated GCase |
| 10 | CA16763 | deglycosylated GCase |
| 11 | CA16764 | deglycosylated GCase |
| 12 | CA16765 | deglycosylated GCase |
| 13 | CA16766 | deglycosylated GCase |
| 14 | CA16767 | deglycosylated GCase |
| 15 | CA16768 | glycosylated GCase |
| 16 | CA16769 | glycosylated GCase |
| 17 | CA16770 | deglycosylated GCase bound to CBE |
| 18 | CA16771 | deglycosylated GCase |
| 19 | CA16772 | glycosylated GCase |
| 20 | CA16773 | glycosylated GCase bound to CBE |

C

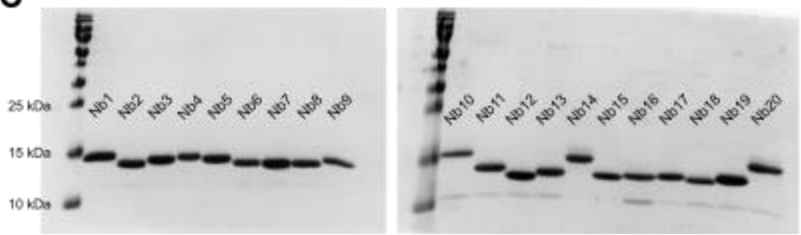


**Supplementary Figure S1: Overview of classification, panning strategy and purification of the full set of 20 Nbs. (A)** CDR3 sequences of the full set of 20 Nbs. **(B)** Conditions in which each Nb has been selected. **(C)** Purification of the full set of 20 Nbs and purity analysis using SDS-PAGE.

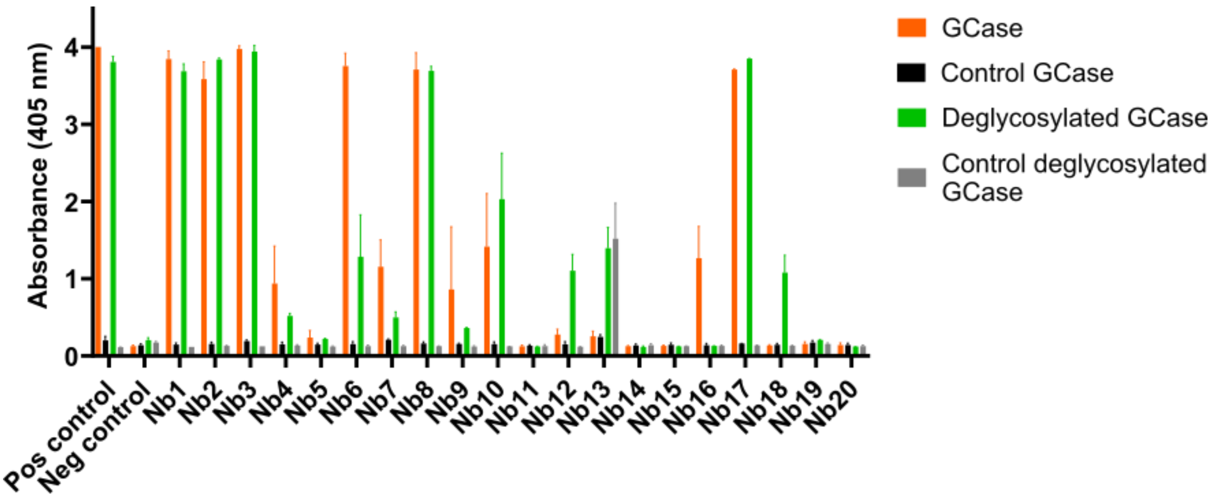


**Supplementary Figure S2: Binding of the set of 20 nanobodies to wild-type and deglycosylated GCase.** ELISA of 20 purified Nbs using GCase or deglycosylated GCase coated on the bottom of the ELISA wells. An irrelevant Nb is used as negative control, while the positive control displays the signal of a Nb (Nb17) directly coated in the ELISA plate. Each ELISA signal is the result of three independent experiments, shown as mean values (bars) with standard deviations (error bars).

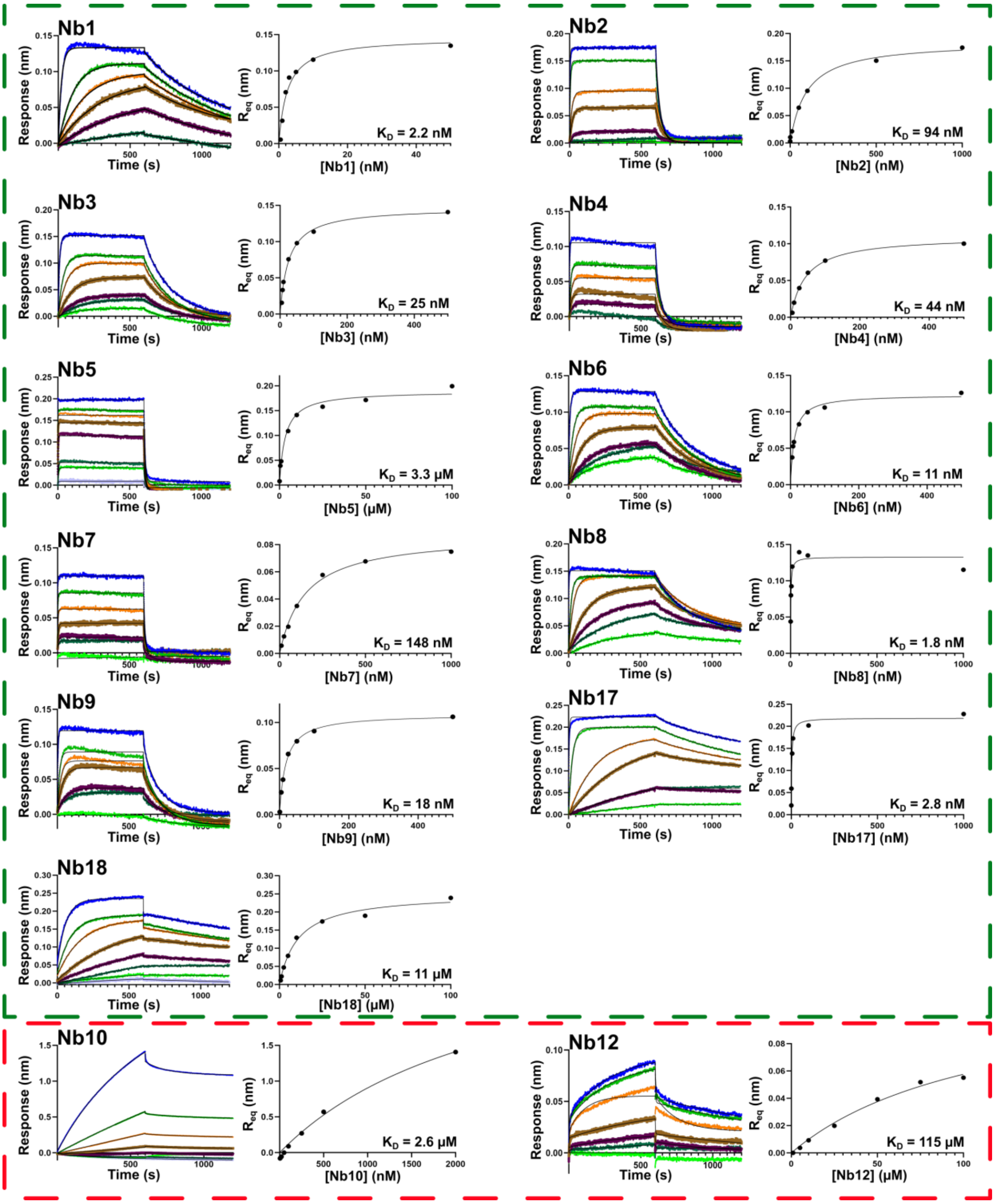


**Supplementary Figure S3: Affinity measurement of the Nbs for GCase using Biolayer Interferometry (BLI).** BLI sensorgrams obtained by titrating increasing concentrations of the Nbs to GCase (green panel) or deglycosylated GCase (red panel) that were trapped on a Streptavidine biosensor, and the most representative resulting dose-response curves, are shown. The sensorgrams were fitted on a 1:1 binding model (ForteBio Analysis Software) and the resulting $R_{eq}$ values were subsequently plotted against the Nb concentration. The $K_D$ values obtained by fitting the dose-response curves with a Langmuir binding equation are given.

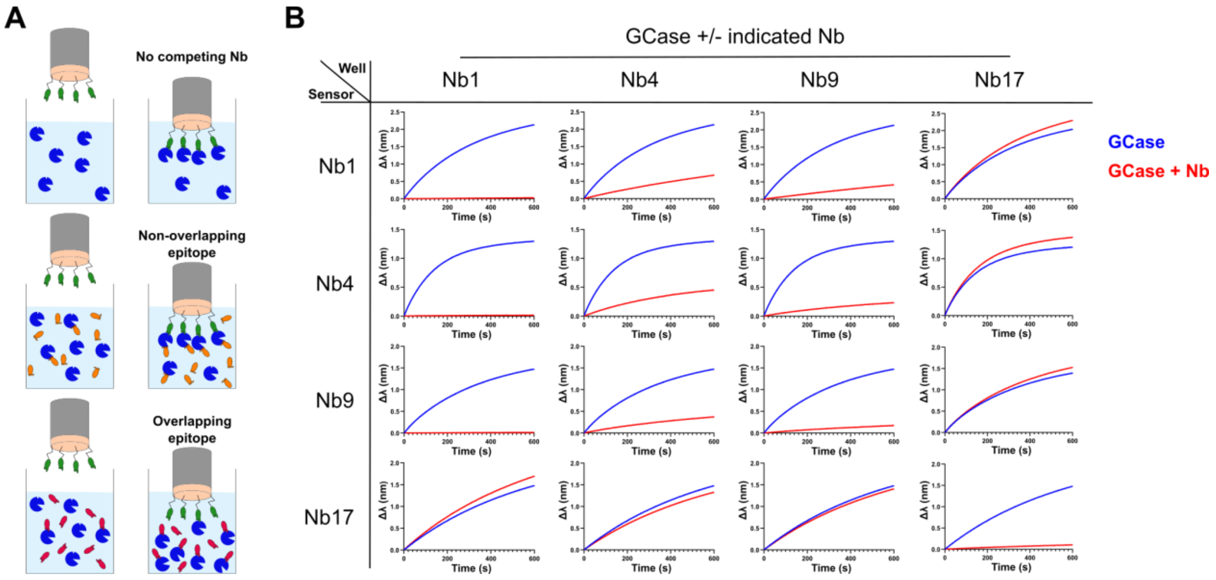


**Supplementary Figure S4: Epitope mapping of a subset of Nbs. (A)** Scheme of the experimental setup. Biotinylated Nbs were loaded on a Streptavidin-coated sensor and then dipped into a solution of GCase in the presence/absence of the different non-biotinylated Nbs. **(B)** Results of the epitope mapping as outlined in panel (A). Blue curves correspond to the binding signal with GCase protein alone, and red curves corresponds to the signal obtained for the same sensor dipped into a GCase+Nb solution.

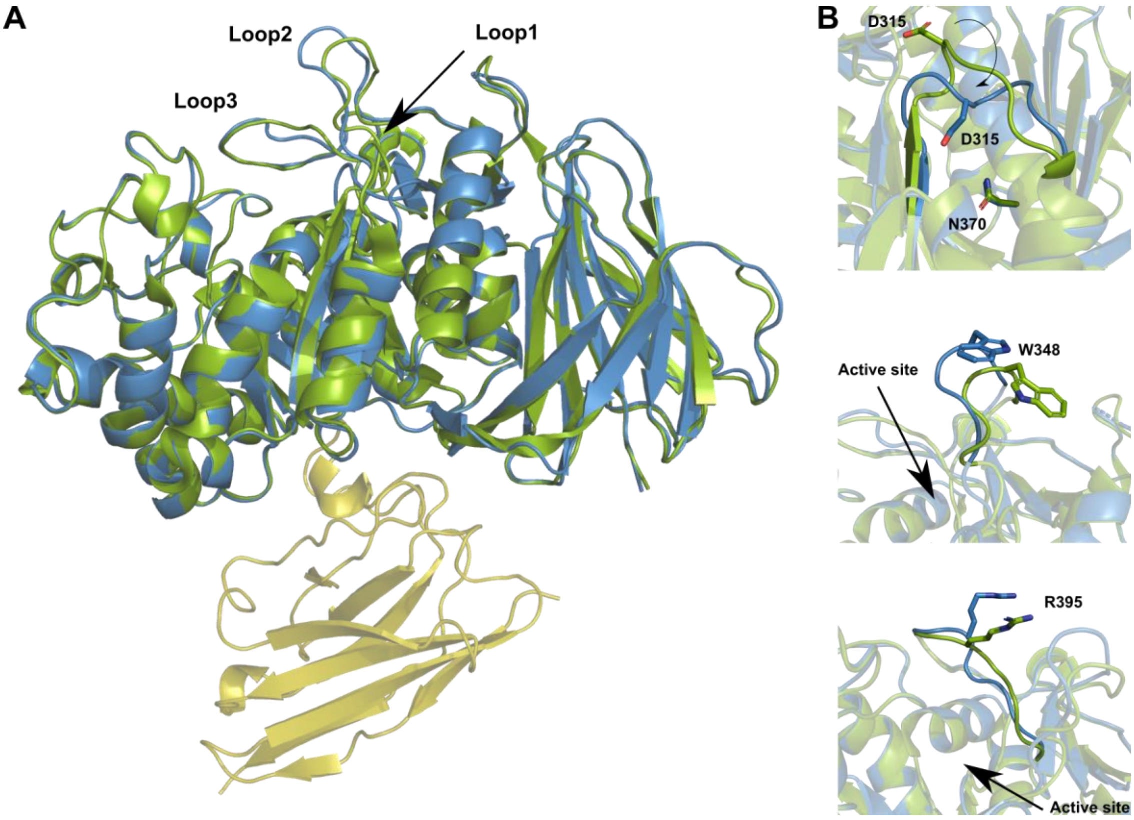


**Supplemental Figure S5: Comparison and conformational changes between GCase bound to Nb1 and unbound GCase (chain A from PDB 1OGS). (A)** Superposition of GCase (blue) from the GCase-Nb1 complex on unbound GCase (green). Nb1 is colored yellow. **(B)** Close-up view of loop1 (*upper panel*), loop2 (*middle panel*) and loop3 (*lower panel*). An arrow indicates the active site, and the most important residues are indicated.

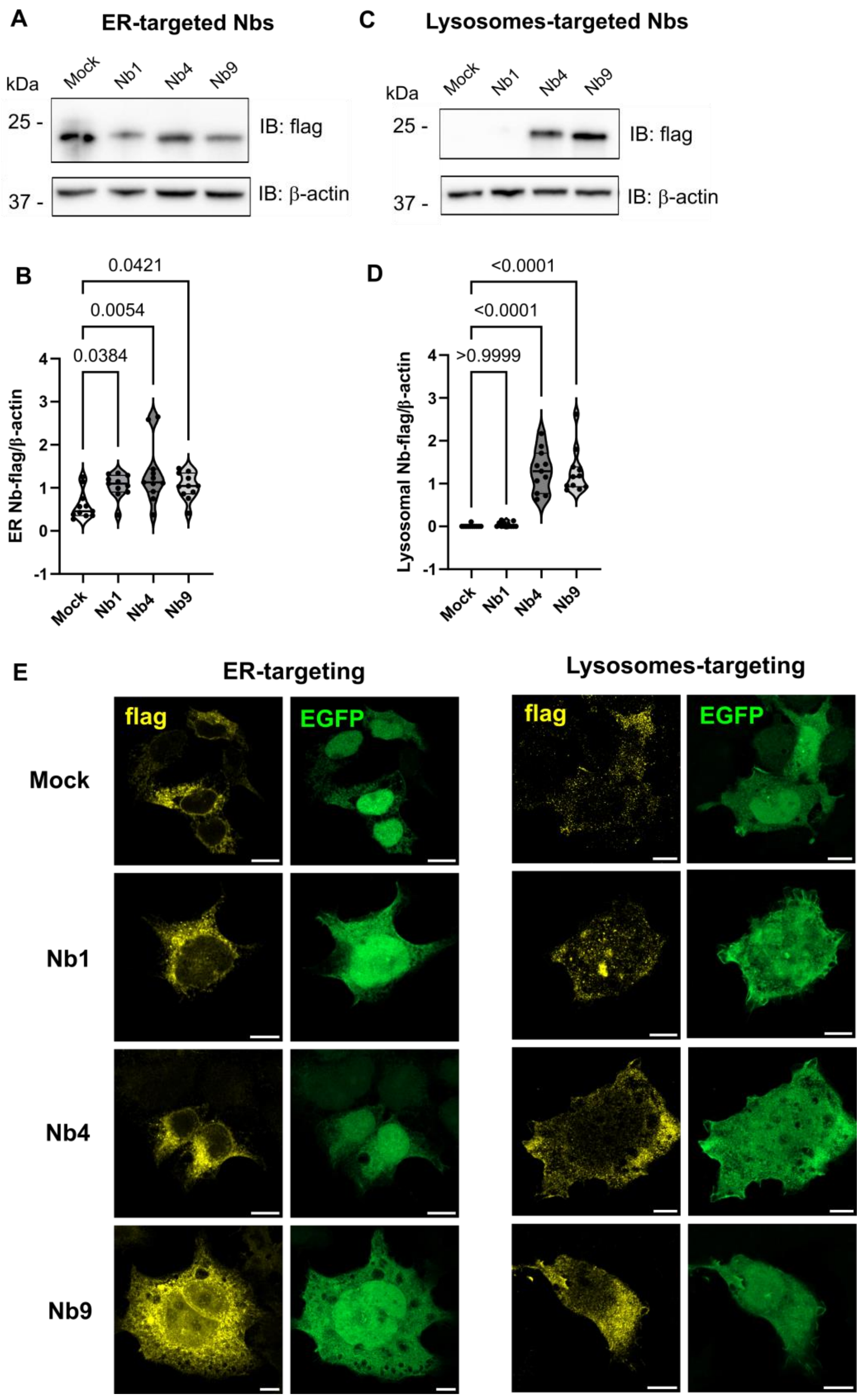


**Supplementary Figure 6: ER- and lysosome-targeted Nbs are expressed differently in the two compartments. (A)** Representative western blot of HEK293T cells transfected with Mock and three selected Nbs targeted to the ER. **(B)** Relative quantification of the flag intensity band, corresponding to the Nbs targeted to the ER, normalized to β-actin shows a variable but consistent expression of all the different Nbs in this compartment (n=9 in 6 independent experiments, data represented as mean±SEM, statistical analysis was performed using a Kruskal-Wallis multiple comparison test). **(C)** Representative western blot of HEK293T cells transfected with Mock and three selected Nbs targeted to the lysosomes. **(D)** Relative quantification of the flag intensity band, corresponding to the Nbs targeted to the lysosomes, normalized to β-actin shows a huge difference in the expression of the different Nbs, with Nb4 and Nb9 presenting the bigger expression (n=11 in 6 independent experiments, data represented as mean±SEM, statistical analysis was performed using a Kruskal-Wallis multiple comparison test). **(E)** Representative confocal images of HEK293T cells overexpressing Mock sequence, Nb1, Nb4 and Nb9 stained with an anti-flag antibody, either targeted to the ER or to the lysosomes, with the relative EGFP fluorescence signal (scale bar 10 µm).

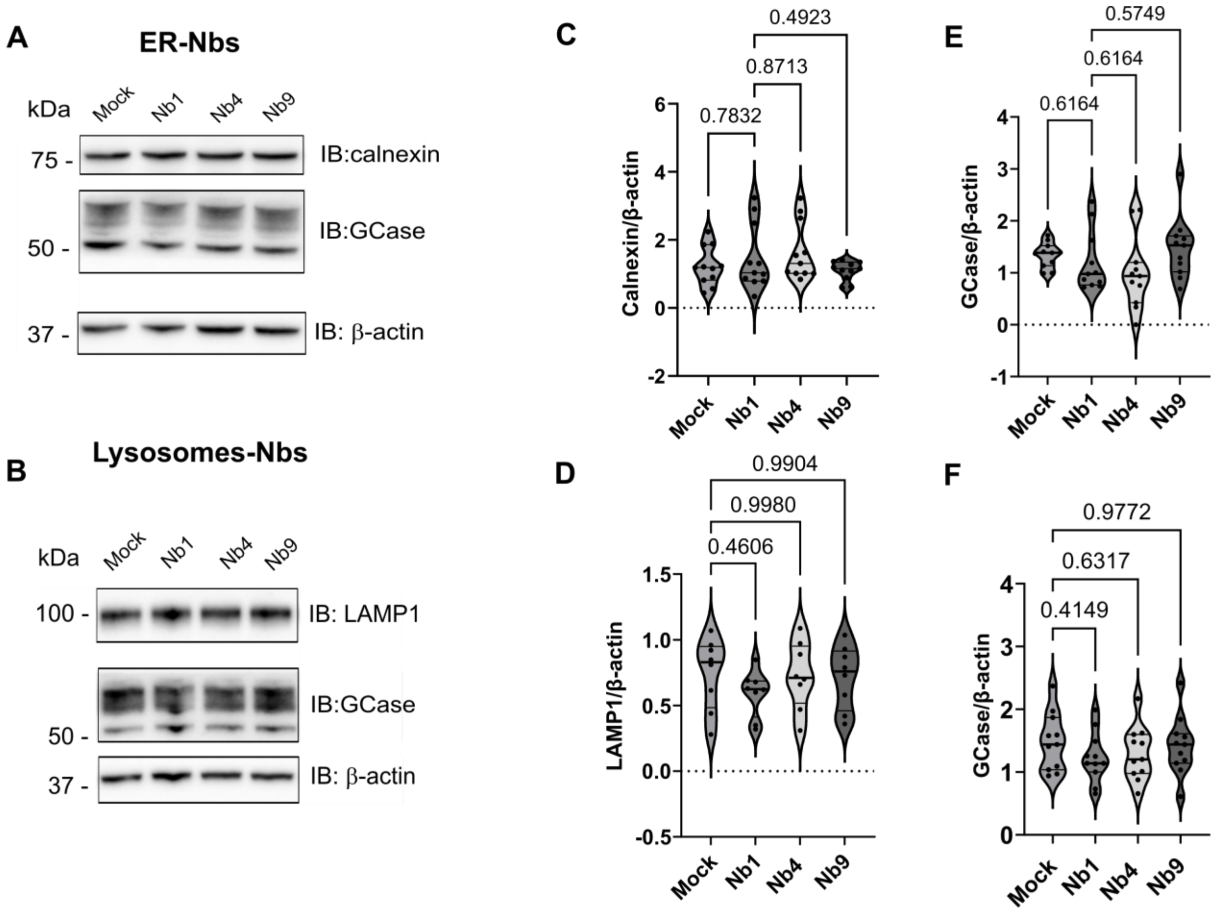


**Supplementary Figure S7: Nb expression does not impact on ER or lysosomal compartments nor on GCase levels. (A)** Representative western blot of HEK293 cells overexpressing the selected ER-Nbs for the GCase and the calnexin levels; **(B)** Representative western blot of HEK293 cells overexpressing the selected lysosome-Nbs for the GCase and the LAMP1 levels ; **(C)** Calnexin expression levels, as quantified by densitometry analysis, show no differences between Mock transfected cells and Nb-expressing cells (n=6-8 in 6 independent experiments, statistical analysis performed via One Way Anova with multiple comparison, DF Nbs 3, DF residual 40, F values 1.305). **(D)** The LAMP1 expression level is unchanged among the different conditions (n=6-8 in 6 independent experiments, data represented as violin plots, statistical analysis performed via One Way Anova with multiple comparison, DF Nbs 3, DF residual 28, F values 0.6592); **(E)** Quantification of GCase expression level in cells expressing the ER-targeted Nbs shows no differences compared to Mock cells (n=9-11, in 6 independent experiments, data represented as violin plots, statistical analysis performed via One Way Anova, DF Nbs = 3, DF = residual 40, F = 1.542); **(F)** Quantification of GCase expression level in cells expressing the lysosomes-targeted Nbs show no differences compared to Mock cells (n=9-11, in 6 independent experiments, statistical analysis performed via One Way Anova, DF Nbs 3, DF residual 40, F values 0.7367).

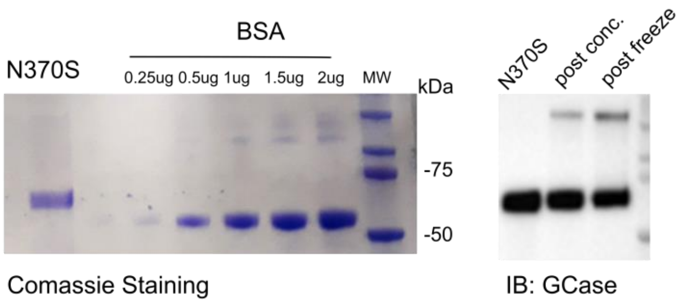


**Supplementary Figure S8: N370S GCase mutant purification from 293T Freestyle cells provided a yield of about 300 µg from 1L of cell culture. (A)** SDS-PAGE analysis quantification using coomassie gel staining and BSA to obtain a standard curve. **(B)** Western blot analysis was performed in order to identify aggregation and degradation using a GCase specific antibody.

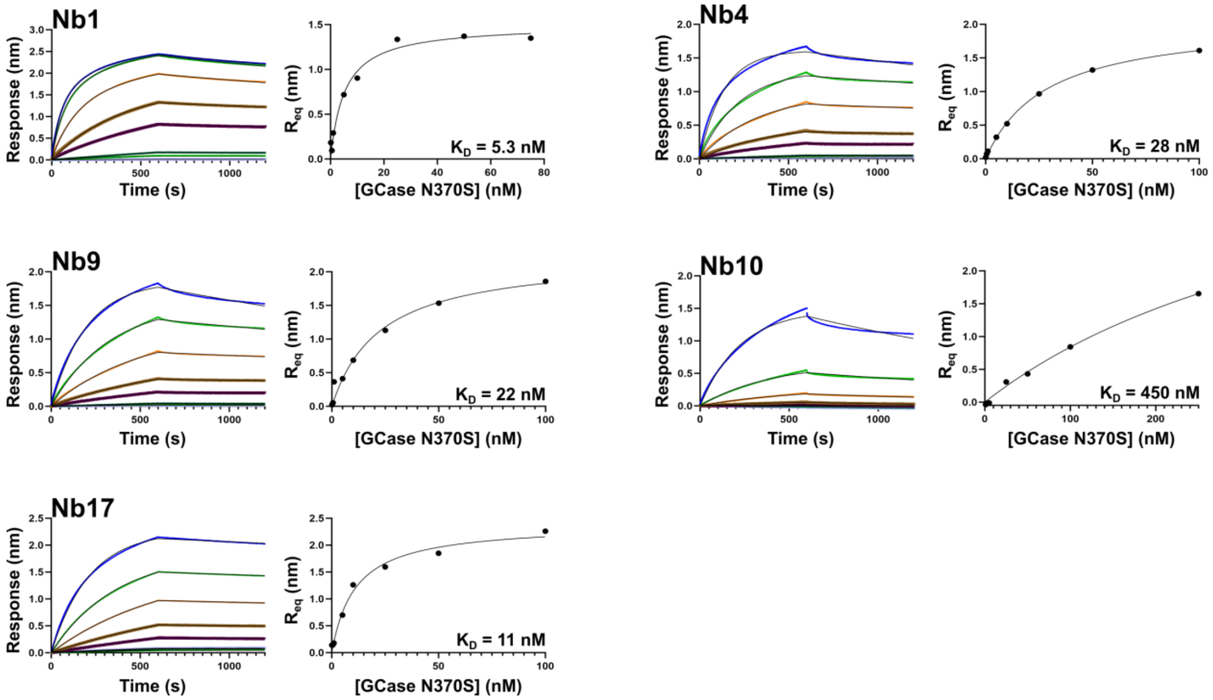


**Supplementary Figure S9: Affinity measurement of the Nbs for GCase N370S using Biolayer Interferometry (BLI).** BLI sensorgrams obtained by titrating increasing concentrations of the GCase N370S to Nbs that were trapped on a Streptavidine biosensor, and the resulting dose-response curves, are shown. The sensorgrams were fitted on a 1:1 binding model (FortéBio Analysis Software), and the resulting $R_{eq}$ values were subsequently plotted against the GCase N370S concentration. The $K_D$ values obtained by fitting the dose-response curves with a Langmuir binding equation are given.

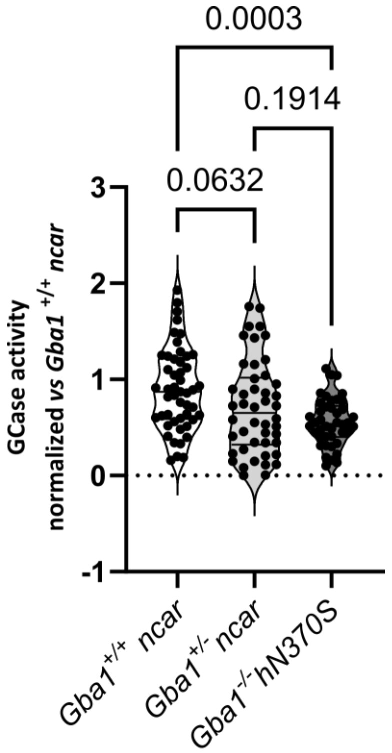


**Supplementary Figure S10: 4-MU GCase activity in gut lysates from *Gba1*$^{+/+}$, *Gba1*$^{+/-}$ and *Gba1*$^{-/-}$ *h*N370S.** *Gba1*$^{-/-}$*h*N370S gut lysates showed a decreased 4-MU GCase activity as compared with tissues from *Gba1*$^{+/+}$ *ncar* animals, while *Gba1*$^{+/-}$ *ncar* tissues show only a trend toward the decrease as compared to wild type tissues (n = 4 experiments, each measuring 4 individual tissues per genotype in three technical replicates, data represented as violin plots, statistical analysis performed via One Way Anova, DF Nbs = 2, DF Residual = 134, F = 8.065).

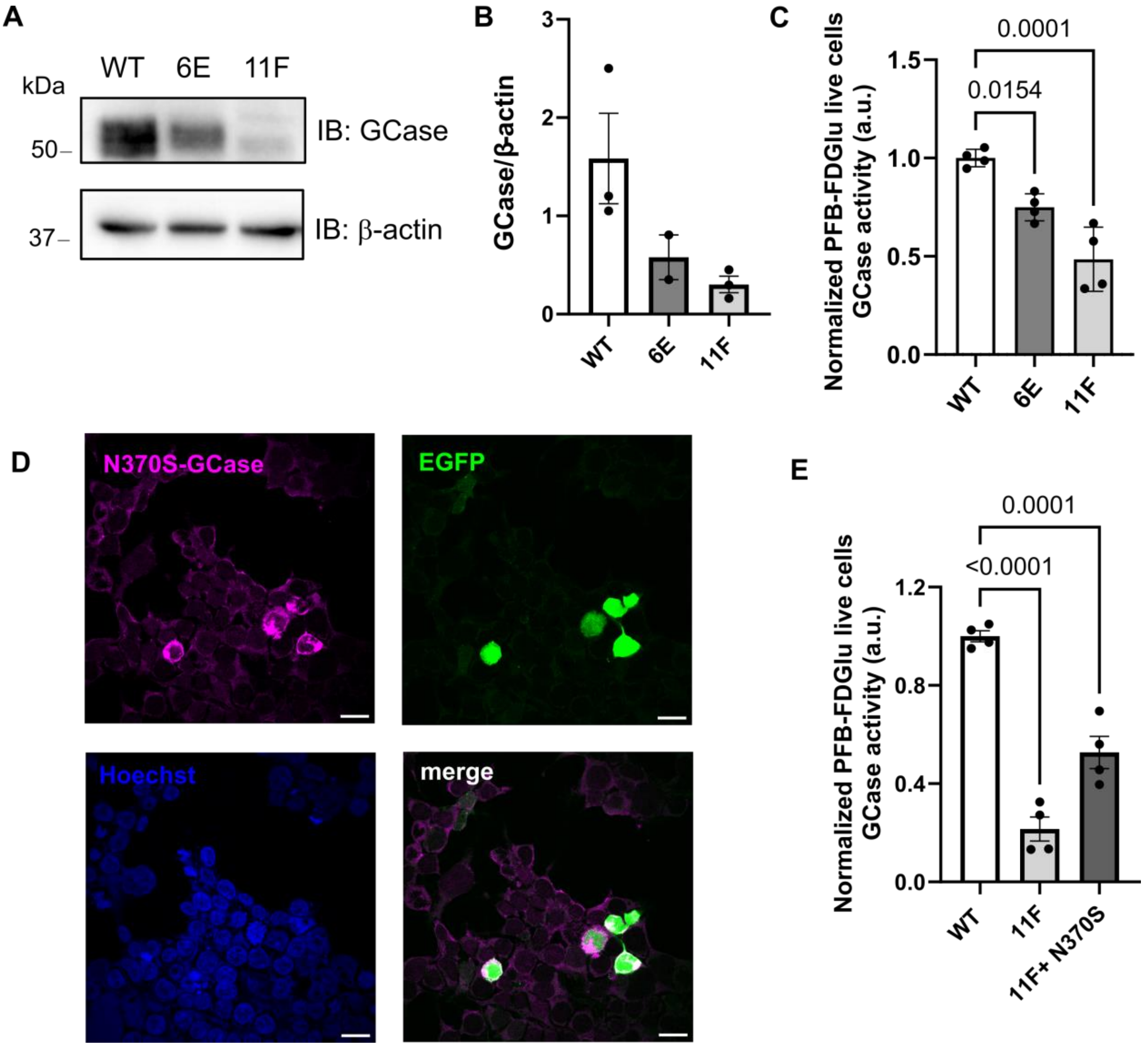


**Supplementary Figure S11: Characterization of *GBA1* knock down HEK293T cells and coexpression of Nbs and N370S GCase mutant. (A)** Representative western blot of 6E and 11F *GBA1* knock down cell clones showing reduced GCase levels as compared with WT and **(B)** relative quantification represented as histograms showing mean±SEM and individual data points (n=2-3 biological replicates measured in independent experiments). **(C)** Lysosomal GCase activity in live cells is significantly reduced in both 6E and 11F *GBA1* knock down cells, as compared with wild type HEK293T cells (n=4 in 2 independent experiments, statistical analysis performed via One Way Anova with multiple comparison, after Shapiro-Wilk test for Normality, DF genotype = 2, DF Residual = 9, F = 23.82). (**D)** Representative image of cotransfection efficiency of Nbs and N370S GCase mutant as evaluated by confocal microscopy imaging (scale bar 10 μm). **(E)** Lysosomal GCase activity in live cells is significantly increased in 11F *GBA1* knock down cells when overexpressing the N370S mutant, as compared with 11F *GBA1* knock down cells (n=4 in 2 independent experiments, statistical analysis performed via One Way Anova with multiple comparison, after Shapiro-Wilk test for Normality, DF genotype = 2, DF Residual = 9, F = 64.75).

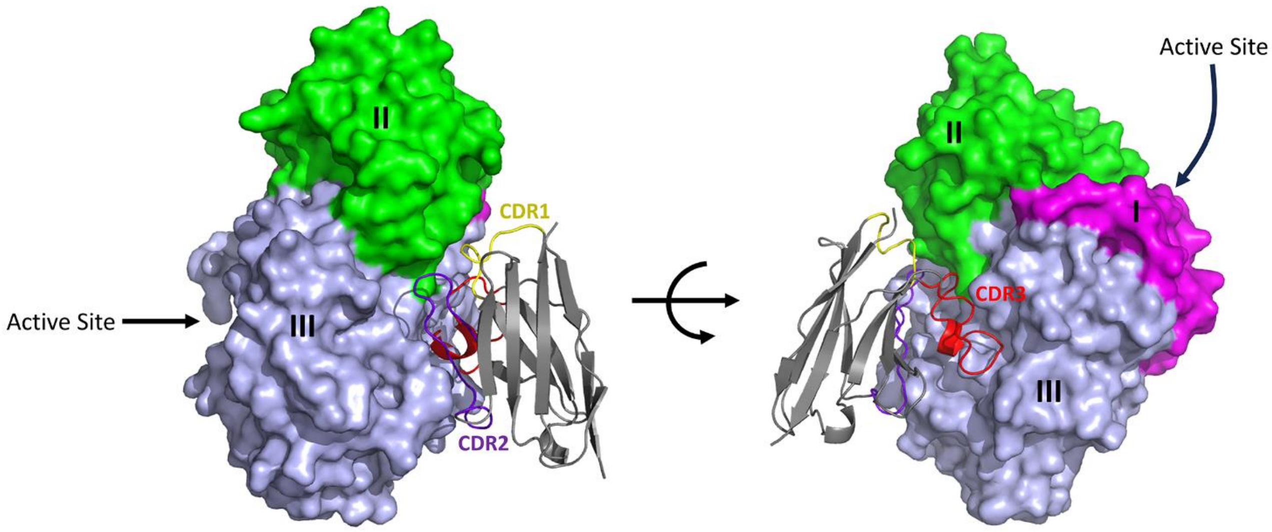


**Supplementary Figure S12: Nb1 binds at the interface of GCase domain II and III and at an opposing side of the GCase active site.** GCase is shown in surface representation with its domains in pink (domain 1), green (domain 2) and light blue (domain 3). Nb1 is colored grey with the CDR1 loop in yellow, CDR2 loop in dark blue and CDR3 loop in red.

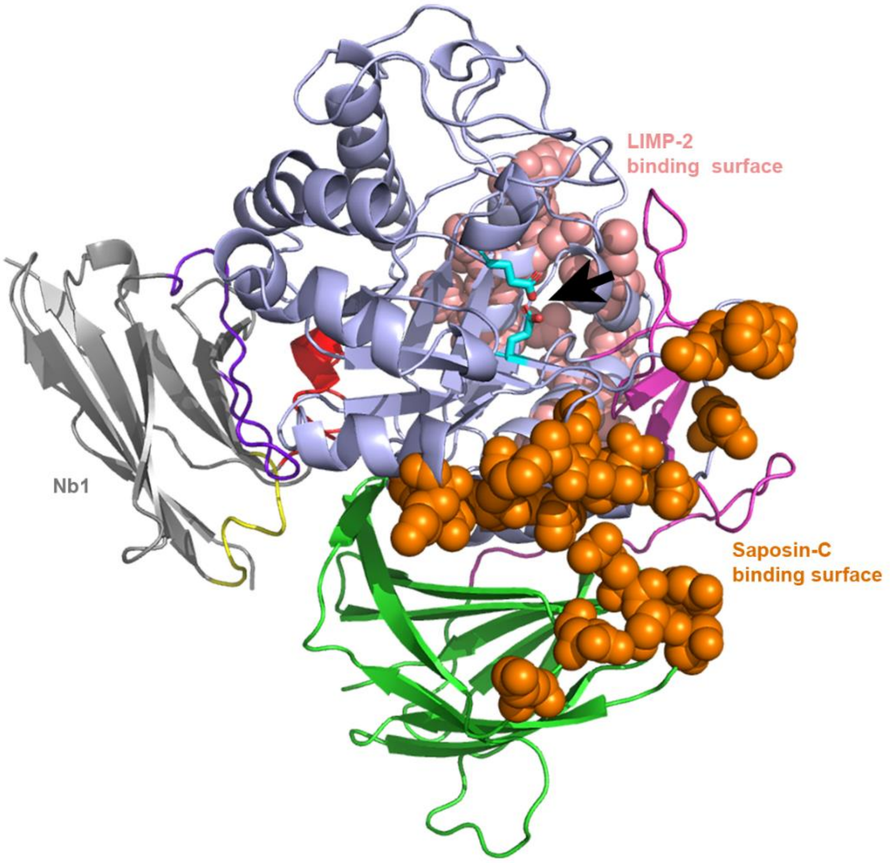


**Supplementary Figure S13: The Nb1 epitope on GCase does not overlap with the presumed Saposin-C and LIMP-2 binding surfaces.** The three domains of GCase are colored in pink (domain 1), green (domain 2) and light blue (domain 3). Residues of GCase that are located on the presumed Saposin-C and LIMP-2 binding surfaces are shown as orange and salmon spheres, respectively[1,2]. The position of the GCase active site is indicated with an arrow. Nb1 is colored grey with the CDR1 loop in yellow, CDR2 loop in dark blue and CDR3 loop in red.

**Supplementary references**


1 Romero R, Ramanathan A, Yuen T, Bhowmik D, Mathew M, Munshi LB *et al.* Mechanism of glucocerebrosidase activation and dysfunction in Gaucher disease unraveled by molecular dynamics and deep learning. *Proc Natl Acad Sci U S A* 2019; **116**: 5086–5095.

2 Zunke F, Andresen L, Wesseler S, Groth J, Arnold P, Rothaug M *et al.* Characterization of the complex formed by β-glucocerebrosidase and the lysosomal integral membrane protein type-2. *Proc Natl Acad Sci* 2016; **113**: 3791–3796.
